# Nanobodies against the myelin enzyme CNPase as tools for structural and functional studies

**DOI:** 10.1101/2024.05.25.595513

**Authors:** Sigurbjörn Markusson, Arne Raasakka, Marcel Schröder, Shama Sograte-Idrissi, Amir Mohammad Rahimi, Ommolbanin Asadpour, Henrike Körner, Dmitri Lodygin, Maria A. Eichel-Vogel, Risha Chowdhury, Aleksi Sutinen, Gopinath Muruganandam, Manasi Iyer, Madeline H. Cooper, Maya K. Weigel, Nicholas Ambiel, Hauke B. Werner, J. Bradley Zuchero, Felipe Opazo, Petri Kursula

## Abstract

2’,3’-cyclic nucleotide 3’-phosphodiesterase (CNPase) is an abundant constituent of central nervous system non-compact myelin, frequently used as a marker antigen for myelinating cells. The catalytic activity of CNPase, the 3’-hydrolysis of 2’,3’-cyclic nucleotides, is well characterised *in vitro*, but the *in vivo* function of CNPase remains unclear. CNPase interacts with the actin cytoskeleton to counteract the developmental closure of cytoplasmic channels that travel through compact myelin; its enzymatic activity may be involved in adenosine metabolism and RNA degradation. We developed a set of high-affinity nanobodies recognizing the phosphodiesterase domain of CNPase, and the crystal structures of each complex show that the five nanobodies have distinct epitopes. One of the nanobodies bound deep into the CNPase active site and acted as an inhibitor. Moreover, the nanobodies were characterised in imaging applications and as intrabodies, expressed in mammalian cells, such as primary oligodendrocytes. Fluorescently labelled nanobodies functioned in imaging of teased nerve fibers and whole brain tissue sections, as well as super-resolution microscopy. These anti-CNPase nanobodies provide new tools for structural and functional biology of myelination, including high-resolution imaging of nerve tissue.

## Introduction

Myelin is a specialised multilamellar membrane that ensheathes the axons of neurons in the central (CNS) and peripheral nervous system (PNS). It plays a crucial role in facilitating rapid and efficient transmission of electrical signals between neurons, enabling the functioning of the vertebrate nervous system [1]. Myelin is formed by specialised glial cells, oligodendrocytes in the CNS and Schwann cells in the PNS. These cells wrap their plasma membrane around the axon, creating a multilayered proteolipid sheath [2, 3]. The myelin sheath is not continuous around a single axon, but rather segmented into internodes, separated by gaps called nodes of Ranvier. Myelin serves as an electrical insulator, preventing leakage of electrical signals along the axon. This allows for rapid and efficient saltatory conduction, in which the electrical signal jumps from node to node, enabling faster transmission of information compared to unmyelinated axons. Myelin also provides structural support to axons, protecting them from damage and degeneration. Additionally, myelin plays a role in nutrient transport and waste removal from axons. Disruptions in myelin formation or maintenance can lead to a variety of neurological disorders, which can range from mild to severe, affecting various aspects of nervous system function. Understanding the biology of myelin and the mechanisms underlying myelin disorders will be crucial for developing effective treatments in the future.

CNPase, or 2’,3’-cyclic nucleotide 3’-phosphodiesterase, is an enzyme of the 2H phosphodiesterase family [4–6], highly expressed in myelinating cells [7, 8]. Deficiency of CNPase in mice causes ultrastructural defects of the axon/myelin-unit [8, 9] and impairs the initiation of executive functions [10, 11]. A homozygous missense mutation in the human gene encoding CNPase correlates with white matter loss and associated neurodegeneration [12], and *CNP* gene mutations in dogs have been linked to lysosomal storage disease and myelin abnormalities [13, 14]. Due to its high abundance in myelinating glia, CNPase is a commonly used marker of myelin, being localised in the non-compacted subcompartments. The molecular function of CNPase is not fully understood, and a high-resolution structure of full-length CNPase has not been experimentally solved. The catalytic function of the N-terminal polynucleotide kinase-like domain is controversial, while the reaction catalysed by the C-terminal phosphodiesterase catalytic domain has been known *in vitro* for >60 years [15] and characterised mechanistically in detail [16–18]. However, it is unclear if the physiological function of CNPase is related to its enzymatic properties, or if it has evolved to be more relevant through its molecular interactions with other proteins [19, 20], RNA [16, 21, 22], and lipid membranes [23, 24]. To characterise different aspects of CNPase function, solve the high-resolution structure of full-length CNPase, and improve the use of CNPase as an imaging and diagnostic tool, new approaches are required.

Nanobodies (Nb), also known as single-domain antibodies or single variable domains on a heavy chain (VHH), are small antigen-binding fragments derived from heavy-chain-only antibodies found in camelids and cartilaginous fishes. Unlike conventional antibodies, which consist of two heavy chains and two light chains, Nbs are composed of a single VHH that has strong, specific antigen-binding capacity. Nanobodies can be produced recombinantly, and their small size and high stability make them suitable for a wide range of applications, from structural to functional studies, as well as in advanced microscopy [25, 26].

We generated and characterised a set of five high-affinity anti-CNPase nanobodies (NbCNP) that can be used in various applications, including structural and functional studies. All NbCNPs were co-crystallised with the catalytic domain of mouse CNPase, allowing the atomic-level identification of five different antigenic epitopes on the CNPase surface. One Nb bound deep into the CNPase active site, acting as an inhibitor, while another one activated CNPase slightly. The high affinity of the NbCNPs allowed further validation in the imaging of nerve tissue using confocal and STED microscopy, and they could be expressed in cultured oligodendrocytes; in addition, their function as intrabodies in living cells was confirmed. The anti-CNPase nanobodies provide state-of-the-art tools for structural biology, as well as high-resolution imaging of myelinated tissue and functional intervention of CNPase-related processes.

## Materials and Methods

### Synaptosome preparation

Wild-type Wistar rats (*Rattus norvegicus*) were obtained from the University Medical Center Göttingen and handled according to the specifications of the University of Göttingen and the local authority, the State of Lower Saxony (Landesamt für Verbraucherschutz, LAVES, Germany). The local authority (Lower Saxony State Office for Consumer Protection and Food Safety) approved the animal experiments (license number: T09/08). Rat brain synaptosomes were enriched as described [27]. Briefly, brains were homogenised in precooled sucrose buffer (320 mM sucrose, 5 mM HEPES, pH 7.4). After centrifugation at 1000 g for 2 min, the supernatant was further centrifuged at 15000 g for 12 min. A discontinuous Ficoll density gradient was applied. The fractions at the interface of 9% Ficoll were pooled, washed in sucrose buffer, and used for immunisation.

### Immunisation

Two alpacas were immunised with enriched rat synaptosomes. The procedure was performed by Preclinics GmbH (Potsdam, Germany). Six injections were performed weekly with 0.5 mg of protein from enriched synaptosomes. Two weeks after the last immunisation, a single boost with 0.5 mg of synaptosomes was performed, and 100 ml of blood was taken 3 and 5 days after the boost immunisation. Peripheral blood mononuclear cells (PBMCs) were isolated using a Ficoll gradient, and the remaining serum was stored at -80 °C. Total RNA was extracted using an RNA extraction kit (Qiagen).

### Enrichment of IgG2 & IgG3 from plasma

Plasma from the two fully immunised animals was enriched in IgG2 and IgG3 following the original protocol [26]. Filtered plasma was injected into a HiTrap protein G HP (Cytiva) column, and the flowthrough was collected and injected into a HiTrap protein A (Cytiva) column. Bound IgGs on the protein G column were eluted first at pH 3.5, followed by a second elution using pH 2.7. The IgG bound to the protein A column was eluted at pH 4.0. Eluted fractions were neutralised using 1 M Tris-HCl, pH 9.0. The fractions were analyzed on denaturing SDS-PAGE, and IgG2 and IgG3 fractions were pooled.

### Plasma-ELISA

The purified full-length human and mouse CNPase (hCNPase and mCNPase) were immobilised overnight at +4 °C on a 96-well immunosorbent plate (Nunc). All the following steps were done by gentle shaking on an orbital shaker. Wells were washed with PBS and blocked with 5% (w/v) skim Milk in PBS for 3 h at RT. After rinsing the wells with PBS, the enriched IgG2-IgG3 mixture was added in a concentration of 0.5 µg/µl and incubated on the wells at RT. Bound IgG2 and IgG3 were then revealed with a monoclonal mouse anti-camelid antibody coupled to HRP (Preclinics, clone: P17Ig12) diluted 1:2000 in PBS. The ELISA was revealed by adding 100 µl of TMB substrate (ThermoScientific) until the blue color was stable. The reaction was quenched with 100 µl of 2 M sulfuric acid. The absorbance was read at 430 nm (BioTek Cytation).

### Nanobody library generation

Total mRNA was extracted from the enriched PBMC from the 2 alpacas using a standard RNA extraction kit (Qiagen). As described [26], the recovered mRNA was retrotranscribed to cDNA using Superscript IV (Invitrogen) and the Cal 0001/2 primers. PCR products were diluted to 5 ng/µl, and 1.5% agarose gel electrophoresis was used to confirm the size of the PCR product. The obtained phagemids were purified using a PCR purification kit (Qiagen). The library was then electroporated into TG1 bacteria. For the transformation, 65 ng of DNA was added to 50 µl of TG1 and repeated 20 times. The electroporated bacteria were left 1 h at 37 °C, then pooled together in 400 ml of 2YT medium (ThermoFisher) supplemented with antibiotics and cultured overnight at +37 °C.

The next day, bacteria were pelleted and resuspended in 25 ml LB medium (ThermoFisher) containing 25% glycerol. The library was aliquoted, snap-frozen in liquid nitrogen, and stored at -80 °C.

### Phage display

A 1-ml library aliquot was grown in 500 ml of 2YT supplemented with antibiotic and grown at +37 °C until OD_600_ reached ∼0.5. Next, ∼1×10^12^ M13KO7 Helper Phages (NEB) were added to the culture, allowing infection for 45 min. Infected bacteria were incubated overnight at +30 °C to produce phages. The next day, the culture supernatant was incubated with 4% (w/v) PEG8000 and left on ice for <2 h to allow phages to precipitate. After several washes in PBS, phages were filtered through a 0.45-µm syringe filter (Sartorius). Full-length human and mouse CNPase were conjugated to desthiobiotin-*N*-hydroxysuccinimide ester (Berry and Associates). 1-3 nmol of mixed antigen was bound to pre-equilibrated Dynabeads MyOne Streptavidin C1 (ThermoFisher). The purified phages were mixed with the beads pre-loaded with a combination of hCNPase and mCNPase and incubated for 2 h at RT. Beads were thoroughly washed in PBS-T. CNPase and bound phages were eluted using 50 mM biotin in PBS. The eluted phages were used to reinfect TG1 cells and initiate another panning cycle. After the last panning round, bacteria were plated on LB agar supplemented with antibiotics, and the following day, 96 colonies were picked and grown in 96-deep-well plates.

### Phage and ramp ELISAs

Each clone in the 96-deep-well plate was infected with helper phages and incubated overnight. Bacteria were centrifuged down, and supernatants containing phages were used directly. The desthio-biotinylated mCNPase and hCNPase were immobilised for 2 h at RT on streptavidin-coated flat-bottom 96-well plates (Thermo Fisher Scientific). The wells were washed three times 10 min with PBS and blocked using 5% skim milk in PBS-T for 3 h at RT. Next, 25 µl of phages from each well were incubated with immobilised hCNPase or mCNPase 2 h at RT. Bound phages were detected by staining for 1 h with anti-major coat protein M13-HRP (Santa Cruz) diluted 1:1000 in 100 µl PBS. TMB substrate was added to each well, and the colorimetric reaction was stopped with 100 µl of 2 M sulfuric acid. The absorbance was read at 430 nm using a plate reader (BioTek Cytation).

### Tissue imaging with anti-CNPase nanobodies

*Nanobody expression and purification for imaging applications –* Nanobodies for imaging were produced in SHuffle® Express (NEB) cells, using a vector coding for a His_14_ tag and bdSUMO as fusion partners [28, 29]. Bacteria were grown in Terrific Broth supplemented with kanamycin at +30 °C. When OD_600_ reached ∼3, 0.4 mM IPTG was added. Induction was allowed for ∼16 h. Cultures were centrifuged and the pellet resuspended in cold lysis buffer (LysB: 100 mM HEPES, 500 mM NaCl, 25 mM imidazole, 2.5 mM MgCl_2_, 10% v/v glycerol, 1 mM DTT, pH 8.0) supplemented with 1 mM PMSF. After disruption by sonication, the lysate was centrifuged at ∼11000 g for 1.5 h at +4 °C. The supernatant was incubated with LysB-equilibrated Ni^+^ beads (cOmplete, Roche) for 1 h at +4 °C. Beads were washed with 3 CV using LysB buffer and with 5 CV of high salt buffer (HSB; 50 mM HEPES, 1.5 M NaCl, 25 mM imidazole, 2.5 mM MgCl_2_, 5% v/v glycerol, 1 mM DTT, pH 7.5). Finally, beads were washed in the buffer of choice for the next application. Elution was carried out using bdSENP1 protease cleavage on the column [28]. Eluted nanobodies were evaluated on SDS-PAGE.

*Fluorophore conjugation –* Purified nanobodies bearing an ectopic cysteine (at their C-terminus or N- and C-termini) were reduced for 1 h on ice using 10 mM tris(2-carboxyethyl)phosphine (TCEP). Excess TCEP was removed using a NAP-5 column (GE Healthcare) pre-equilibrated with cold-degassed PBS, pH 7.4. Freshly reduced nanobodies were immediately mixed with ∼3-5-fold molar excess of maleimide-functionalised fluorophore and incubated for 2 h at RT. Excess dye was removed using a Superdex™ 75 increase 10/300 GL column (Cytiva) on the Äkta-Prime FPLC system.

*Histology and immunohistochemistry –* Naïve C57BL/6 mice were intracardially perfused with saline (5 min) followed by a fixative containing 4% PFA (10 min). Samples were post-fixed in the same fixative for 24 h, transferred into 30% sucrose, and embedded in OCT. CNPase staining was performed on frozen sections of mouse brain. Tissue sections were first blocked and permeabilised with 10% (v/v) FCS and 0.2% (v/v) Tween-20 for 30 min at RT. The following fluorescently labelled nanobodies were used for overnight incubation at +4 °C: Nb5E-Star635p, Nb8D-Star635p, and Nb10E-Star635p. Sections were washed in PBS/0.05% (v/v) Tween-20 and stained with DAPI. Images were acquired with a VS120-L100-J slide scanner (Olympus) using a 10x objective.

*Nerve preparation –* For immunolabelling of teased fiber preparations, sciatic nerves dissected from mice were transferred into ice-cold PBS and processed as described [30]. Mice were bred and kept in the mouse facility of the Max Planck Institute of Multidisciplinary Sciences registered according to §11 Abs. 1 TierSchG and sacrificed in accordance with the German animal protection law (TierSchG) and approved by the Niedersächsisches Landesamt für Verbraucherschutz und Lebensmittelsicherheit (LAVES) under license 33.19-42502-04-17/2409. Briefly, using two fine forceps (Dumont No. 5), the epineurium was removed from the dissected sciatic nerves, and small nerve pieces were transferred onto a new coverslip. By pulling the fiber bundles carefully apart with both forceps, the axons were separated from each other. Slides were dried and stored at −20 °C for later immunolabelling.

*Immunofluorescence -* Samples were blocked and permeabilised with 3% (w/v) BSA, 0.1% (v/v) Triton X-100 for 20 min at RT with gentle shaking. Antibodies and nanobodies were applied in PBS supplemented with 1.5% BSA and 0.05% Triton X-100 for 1 h at RT with gentle shaking. When indirect detection was needed, secondary antibodies were added for 1 h at RT, followed by several PBS washes and DAPI staining. Coverslips were rinsed in distilled water and mounted using Mowiöl (12 ml of 0.2 M Tris buffer pH 7.5, 6 ml distilled water, 6 g glycerol, 2.4 g Mowiöl 4-88, Merck Millipore). Samples were imaged immediately or within the next 48 h, and samples were kept at +4 °C.

*Confocal & STED microscopy –* Images from mounted samples in Mowiöl were acquired using a STED Expert Line microscope (Abberior Instruments, Göttingen, Germany). The microscopy setup comprised an IX83 inverted microscope (Olympus, Hamburg, Germany) equipped with UPLSAPO 100x 1.4 NA oil immersion objective (Olympus). In addition, 488 nm, 561 nm, and 640 nm lasers were used for confocal imaging. High-resolution images were obtained using the 775 nm pulsed STED depletion laser from the same setup. Images were analyzed in Fiji/ImageJ (v. 1.53o).

### Recombinant CNPase production

Recombinant mCNPase variants (full-length and catalytic domain) and full-length hCNPase were expressed using pTH27 vectors [31] using autoinduction [32] in *E. coli* Rosetta(DE3) and purified as described [33]. The purity of the main SEC fractions was analyzed via SDS-PAGE, and the pure fractions were pooled and concentrated to 18-29 mg/ml. Full-length CNPase was split into 50 μl aliquots, snap-frozen in liquid N_2,_ and stored at -80 °C. The purified mCNPase catalytic domain was stored on ice.

### Anti-CNPase Nb production for structural studies

For large-scale expression and purification for structural studies, NbCNP cDNAs were cloned into pTH27 (N-terminal His_6_ tag; [31]) and the pHMGWA vector (N-terminal His_6_-maltose binding protein (MaBP); [34]), using the Gateway system (Invitrogen).

Nb-CNP 10E was expressed from pTH27 and purified like described for anti-Arc Nbs [35]. TEV proteolysis was carried out for the His-tagged protein after the SEC step, at +37 °C for 2-4 h, and a reverse NiNTA affinity purification step was done to remove TEV protease and the cleaved tag. Final purification was carried out on a Superdex 75 10/300 Increase GL SEC column in 20 mM Tris-HCl pH 7.4, 150 mM NaCl. Fractions assessed pure *via* SDS-PAGE were pooled, concentrated in a 10-kDa MWCO spin concentrator, split into 50 μl fractions, snap-frozen in liquid N_2_, and stored at -80 °C.

Due to low yields, the other NbCNPs were cloned into the pHMGWA vector, resulting in N-terminal His_6_-MaBP fusion constructs. Competent *E. coli* Shuffle T7 cells (New England Biolabs, MA, USA) were transformed with the pHMGWA-MaBP-Nb constructs. A single transformed colony was used to inoculate 10 ml of LB starter culture containing 100 μg/ml ampicillin and incubated overnight at +30 °C and 200 rpm. The starter cultures were diluted 100-fold into 500 ml of the same medium. The cultures were incubated at +30 °C and 200 rpm until OD_600_ reached 0.5-0.7. Protein expression was induced with 0.4 mM IPTG at +20 °C for 20 h. Cells were harvested via centrifugation at 6000 g and +4 °C for 1 h, supernatant discarded and pellets resuspended in 50 mM HEPES, 500 mM NaCl, 5 mM MgCl_2_, 25 mM imidazole, 10% (v/v) glycerol and 1 mM DTT, pH 8.0 (35 ml per 500 ml expression culture) supplemented with 0.1 mg/ml lysozyme and cOmplete EDTA-free protease inhibitors. Cells were lysed via a single freeze-thaw cycle followed by sonication, and the soluble fraction was collected via centrifugation at 30000 g and +4 °C for 1 h. The soluble fraction was filtered through 0.45 μm syringe filters and applied to a NiNTA agarose resin equilibrated in 50 mM HEPES, 500 mM NaCl, 5 mM MgCl_2_, 10% (v/v) glycerol and 1 mM DTT, pH 8.0 and the resin washed with 12 CV of the same buffer. Bound protein was eluted in 5 CV (10 ml) of the same buffer containing 500 mM imidazole. TEV protease was added to the eluate, followed by dialysis against 1 l of 50 mM HEPES, 500 mM NaCl, 5 mM MgCl_2_, 10% (v/v) glycerol, and 1 mM DTT (pH 8.0). As these constructs cleaved poorly, at least 3 mg of TEV protease were added to each 10 ml eluate, and dialysis was maintained at +4 °C for 35 h, renewing the DTT in the buffer after the first 24 h. The proteins were then subjected to reverse NiNTA affinity purification to remove TEV protease, uncleaved fusion protein, and free affinity tag. The flowthrough and wash fractions were concentrated to 1 ml in 10 kDa MWCO spin concentrators and further purified on a Superdex 75 Increase 10/300 GL SEC column in 20 mM HEPES, 150 mM NaCl, 0.5 mM TCEP, pH 7.5. Fractions determined to contain pure Nb by SDS-PAGE were pooled, concentrated to 5-25 mg/ml, and split into 50-μl aliquots before snap-freezing in liquid N_2_ and storing at -80 °C. The identity of all purified nanobodies was confirmed by mass spectrometry. For structural work on nanobody 8D, a protein batch from the His_14_-bdSUMO fusion system (see above) was used, due to high yields of pure protein.

### Folding and stability assays

Differential scanning fluorimetry (DSF), or Thermofluor [36], was used to assess the thermal stability of CNPase constructs upon Nb binding. The assay buffer was 20 mM HEPES, 150 mM NaCl, 0.5 mM TCEP, pH 7.5. Proteins were diluted to 0.5-2 mg/ml in the assay buffer and mixed with 100x SYPRO-Orange (in 50% (v/v) DMSO/assay buffer) in 384-well PCR plates to a final concentration of 5x SYPRO-Orange, making the final DMSO concentration in the assay 2.5% (v/v). The assay volume was 18 μl. Fluorescence emission at 610 nm, following excitation at 465 nm, was measured in a LightCycler 480 LC RT-PCR system (Roche, Basel, Switzerland) over the temperature range 20-95 °C, with a temperature ramp of +2.4°C/min. T_m_ was determined as the maximum of the first derivative of the melting curve. If more than one prominent peak appeared in the first derivative plot, the identity of the main and secondary peaks was determined by the approximate area under the peak.

For nanobody 8D, the stabilisation of CNPase by the Nb was additionally studied with nanoDSF using a Prometheus NT.48 instrument (NanoTemper Technologies, Munich, Germany). NanoDSF is a label-free method based on Trp fluorescence, and nanoDSF samples were at 2 mg/ml in SEC buffer (20 mM HEPES, 200 mM NaCl, 0.5 mM TCEP, pH 7.5). The samples included the nanobody, CNPase, and the complex prepared by SEC.

### Pull-down binding assay

For assessment of Nb binding and crude epitope mapping, 0.5 mg/ml of mCNPase catalytic domain were mixed with an equimolar amount of Nb and incubated on ice for 20-45 min. 200 μl were loaded onto 100 μl of NiNTA agarose resin equilibrated in 20 mM HEPES, 150 mM NaCl, 20 mM imidazole, 0.5 mM TCEP, pH 7.5, and the mixture was incubated at +4 °C under gentle agitation for 1 h, followed by centrifugation at 200 g, +4 °C for 5 min. The supernatant (unbound protein) was decanted and the resin washed three times in the same buffer. To elute bound protein, the resin was incubated in the same buffer with 300 mM imidazole for 30 min before centrifugation as above. The fractions were analysed via SDS-PAGE. His_6_-tagged NbCNP-10E was subjected to pull downs with untagged CNPase (full-length and catalytic domain with and without C-terminal tail) in the same manner.

### Calorimetry

The thermodynamics and affinity of Nb binding to CNPase were measured on a MicroCal iTC200 instrument (Malvern Panalytical, Malvern, UK). Binding of NbCNPs to mCNPase catalytic domain was measured in 20 mM HEPES, 150 mM NaCl, 0.5 mM TCEP, pH 7.5 with 8.5-11 μM CNPase in the cell and 88-95 μM Nb in the syringe at +25 °C, with a reference power of 5 μcal/s and the same injection volumes and timing as above. Data analysis was carried out in Origin. Binding enthalpy (ΔH), association/dissociation constant (K_a_/K_d_), and binding entropy (ΔS) were obtained through fitting to a 1:1 binding model.

### Crystallisation and structure determination

The CNPase catalytic domain was co-crystallised with each of the five NbCNPs using sitting-drop vapour diffusion. Table 1 contains the specific methods for each complex. Diffraction data for the CNPase-Nb complexes were collected using synchrotron radiation on the beamline P11 at PETRAIII/DESY (Hamburg, Germany) [37, 38] at an X-ray wavelength of 1.033 Å on a Dectris Eiger 16M detector and processed using XDS [39] (Table 2). As the data for the NbCNP-5E complex showed moderate anisotropy, anisotropic scaling was carried out for this dataset using STARANISO (http://staraniso.globalphasing.org/cgi-bin/staraniso.cgi) [40]. Data quality was analysed in XTRIAGE [41] and molecular replacement done using PHASER [42].The search model for CNPase was PDB entry 2XMI [16] and for each nanobody, the search model was a close sequence homologue (Table 1). Refinement was carried out in PHENIX.REFINE [43] and manual building in Coot [44]. Structure validation was performed using MolProbity [45]. Table 1 lists specific details of structure determination for each complex, and Table 2 has the data processing and refinement statistics.

**Table 1.**
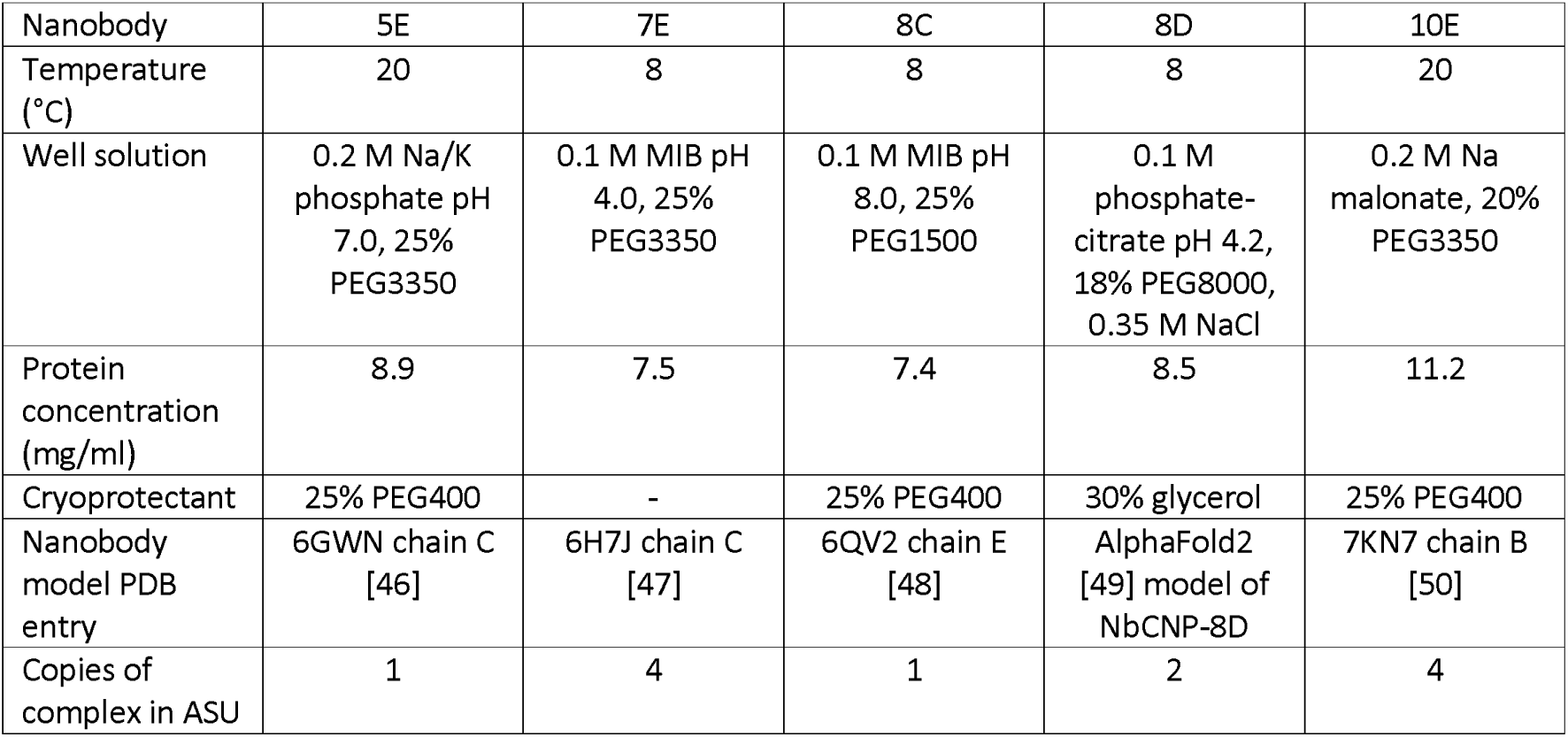
Crystallisation and structure solution of CNPase-nanobody complexes. MIB; buffer mixture of malonic acid, imidazole, and boric acid.

**Table 2.**
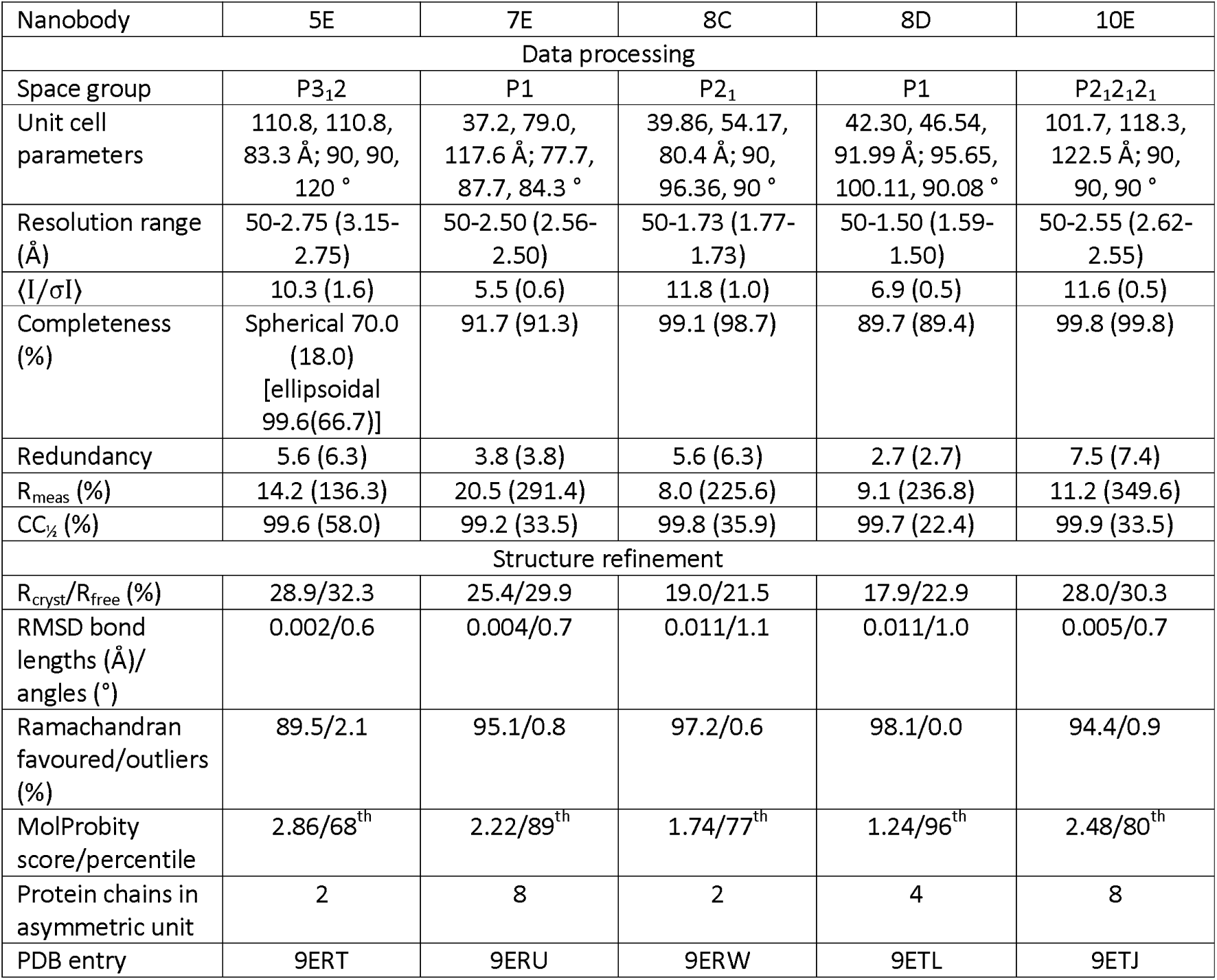
X-ray diffraction data processing and crystal structure refinement. The values in parentheses correspond to the highest-resolution shell.

### Small-angle X-ray scattering (SAXS)

SAXS data were collected on the beamlines SWING [51] at the SOLEIL synchrotron (Gif-sur-Yvette, France) and BM29 [52] at the ESRF (Grenoble, France). To measure solution scattering of CNPase-Nb complexes, Nbs were mixed with the CNPase catalytic domain in 1.3-fold molar excess, and data were collected using a SEC-SAXS setup. The column used was Bio-SEC-3 130 Å 4.6/300 (Agilent Technologies). All samples were run in 20 mM HEPES, 150 mM NaCl, 0.5 mM TCEP, pH 7.5. Data were processed using FOXTROT (SOLEIL synchrotron) and ATSAS [53]. Frame selection and buffer subtraction were carried out in CHROMIXS [54], primary analysis in PRIMUS [55], and distance distribution analysis in GNOM [56]. *Ab initio* models were created using DAMMIN [57] and GASBOR [58] and multi-phase models in MONSA [57]. Theoretical scattering curves for crystal structures were calculated using CRYSOL [59].

### Activity assay *in vitro*

3’-phosphodiesterase activity was measured as described [17, 60], using 2’,3’-cyclic NADP^+^ (Biolog Life Science Institute, Bremen, Germany) as the substrate in a coupled assay. Activity was measured in 100 mM Bis-Tris pH 6.0, 10 mM MgCl_2_ in the presence of 1 U G6P dehydrogenase, 10 mM G6P and 0-2 mM 2’,3’-cNADP^+^ at +25 °C in 96-well plates in a Spark 20M multimode plate reader (Tecan Life Sciences, Switzerland). To initiate the reaction, 2x solutions of serially diluted substrate were mixed with a 2x solution of the reaction buffer, CNPase, G6P, and G6P-dehydrogenase, and the formation of NADPH was monitored for 20 min. Substrate concentrations were determined via absorbance at 259 nm (ε=18 mM^-1^cm^-1^). Initial velocities were plotted as a function of substrate concentration and kinetic parameters obtained by fitting with the non-linear Michaelis-Menten function in GraphPad Prism.

### Expression of anti-CNPase nanobodies as intrabodies

*Construct preparation –* Mammalian expression plasmids encoding fluorescent nanobody constructs were created using InFusion cloning (Takara Bio). DNA fragments encoding the open reading frame (ORF) of the nanobodies with 15-bp complementary overhangs were amplified using PCR and purified from agarose gel bands (Macherey-Nagel Ref 740609.50). A pAAV vector, containing a CMV promoter, the open-reading frame of mRuby3 [61], and a hGH polyA sequence, was linearised using *Age*I-HF and *Xcm*I (New England Biolabs) digestion and purified from an agarose gel band. Linear NbCNP DNA fragments were mixed with the linearised vector and cloned in InFusion reactions, followed by transformation to OneShot Stabl3 chemically competent cells (Fisher Scientific C737303), plasmid propagation and purification, and DNA sequencing (Sequetech Corporation, Mountain View, CA). The obtained constructs had the architecture CMV-NbCNP-mRuby3-hGH polyA, where the nanobody and mRuby3 are linked by the amino acid sequence GDPPVAT. The constructs were amplified in OneShot Stabl3 cells and purified using a plasmid midi kit (Qiagen 12945) for high yields of endotoxin-free DNA. Tom70-EGFP-CNPase-ALFAtag was ordered as a synthetic gene (GeneArt Thermo Scientific) and cloned into the mammalian expression plasmid pcDNA3.1 using Gibson assembly (NEB). Constructs were validated by Sanger sequencing.

*Culture and transformation COS-7 cells –* COS-7 fibroblasts were obtained from the Leibniz Institute DSMZ—German Collection of Microorganisms and Cell Culture (DSMZ Braunschweig, Germany) and cultured in Dulbecco’s MEM supplemented with 10% FBS, 4 mM L-glutamine, 0.6% penicillin and streptomycin, at +37 °C, 5% CO2 in a humified incubator. For immunostaining, cells were plated on poly-L-lysine -coated coverslips in 12-well plates. Cells on coverslips were transfected and co-transfected using 500 ng of each plasmid mixed with 2 µl of Lipofectamine 2000 (ThermoFisher) per coverslip. Cells were fixed using 4% PFA + 0.025% glutaraldehyde ∼16 h after being transfected. Fixed cells were stained with DAPI and mounted on Mowiöl (12 ml of 0.2 M Tris buffer, 6 ml distilled water, 6 g glycerol, 2.4 g Mowiol 4-88, Merck Millipore) for imaging.

*Oligodendrocyte progenitor cell purification –* Oligodendrocyte progenitor cells (OPCs) were purified using an immunopanning protocol from P5-P7 Sprague Dawley rat brains as described [62]. The involved animal procedures were approved by the Institutional Administrative Panel on Laboratory Animal Care (APLAC) of Stanford University and followed the National Institutes of Health guidelines under animal protocol APLAC 32260. The freshly purified primary OPCs were plated onto Ø10-cm tissue culture plates (Fisher Scientific 08-772E; coated with 10 µg/ml poly-D-lysine (PDL), Sigma-Aldrich P6407) in OPC proliferation media (DMEM-Sato) supplemented with 10 ng/ml recombinant human ciliary neurotropic factor (CNTF; Peprotech 450-13), 10 µM forskolin (Sigma-Aldrich F6886), 10 ng/ml recombinant human platelet-derived growth factor AA (PDGF; Peprotech 100-13A) and 1 ng/ml recombinant human neutrophin 3 (NT-3; Peprotech 450-03) and allowed to recover and proliferate at +37 °C/10% CO_2_. Half of the media was replenished every two days, and cell morphology and density were monitored daily until transfection.

*Transfection and differentiation of oligodendrocyte progenitor cells –* Proliferating OPCs were detached from plates using 0.0025% trypsin (Gibco 25300120; diluted using Earle’s balanced salt solution, Sigma-Aldrich E6267) and 30% fetal bovine serum (Gibco 10437028; diluted in Dulbecco’s phosphate-buffered saline (DPBS), Cytiva HyClone #SH30264.01), and collected *via* centrifugation at 90 g for 10 min. For transfection, OPCs were gently resuspended in P3 Primary Cell Nucleofector solution (Lonza Cat. No. V4XP-3032) at 10000 – 12500 cells/µl. For each transfection reaction, 20 µl of cell suspension was mixed with 400 ng of endotoxin-free plasmid DNA. Electroporation was performed using a Lonza 4D-Nucleofector X Unit (AAF-1003X) assembled with a 4D-Nucleofector Core Unit (AAF-1002B) with pulse code DC-218. The electroporated cells were allowed to rest for 10 min at ambient temperature, after which 80 µl of antibiotic-free DMEM-Sato base media was added to each electroporated cell suspension. Each diluted suspension was carefully mixed and further diluted to 2 ml using OPC differentiation media (antibiotic-free DMEM-Sato supplemented with 10 ng/ml CNTF (Peprotech 450-13), 10 µM forskolin (Sigma-Aldrich F6886) and 40 ng/ml 3,31,5-triiodo-L-thyronine (Sigma-Aldrich T6397)). From each suspension, 100 µl aliquots were transferred onto Ø12-mm glass coverslips (coated with 10 µg/ml PDL in 1.5 mM boric acid, pH 8.4) in a 24-well tissue culture plate (Falcon #353047). The cells were allowed to adhere onto the surface for 30 min at ambient temperature, after which 400 µl of OPC differentiation media was added to each well. The cells were differentiated for 5 days at +37 °C/10% CO_2_, with half of the media replenished in each well on day 3 post-transfection.

*Antibodies and staining reagents –* Recombinant rat anti-myelin basic protein (MBP) (Abcam ab7349, 1:20000 dilution) and mouse anti-CNPase (Millipore MAB326R, 1:100 dilution) were used for primary staining of cultured cells. The following secondary antibodies were used at a 1:1000 dilution for cultured cells: donkey anti-rat Alexa Fluor 488 (Invitrogen A-21208) and donkey anti-mouse Alexa Fluor 488 (Invitrogen A-21202). Filamentous actin was stained using phalloidin AlexaFluor 488 conjugate (Invitrogen A12379) at a 7:1000 dilution. To distinguish intact cells from cell debris, HCS CellMask Blue (Thermo Scientific H32720) staining was used at 1:1000 dilution. Rabbit polyclonal anti-NaV1.6 (Alomone labs, #ASC-009) was used in 1:100 dilution followed by 1:500 dilution of a secondary nanobody fused to AberriorStar 580 (NanoTag Biotechnologies, N2402) in the STED experiment.

*Cultured cell staining and epifluorescence microscopy –* On day 5 of differentiation, cell medium was removed by suction, and cells were fixed in 4% paraformaldehyde in PBS for 15 min, followed by three PBS washes. Cells were permeabilised using 0.1% Triton X-100 in PBS for 3 min, followed by three PBS washes. Before primary staining, cells were incubated for 30-60 min in 3% BSA in PBS. All procedures were performed at RT.

For primary staining, diluted primary antibodies in 3% BSA in PBS were incubated overnight at +4 °C. The next day, the primary antibody solutions were removed, followed by three PBS washes and the addition of diluted AlexaFluor 488-conjugated secondary antibodies (anti-rat for MBP and anti-mouse for CNPase staining). Secondary staining was performed for 1 h at RT, followed by three PBS washes. All cells were finally stained using diluted HCS CellMask Blue, followed by three PBS washes.

For actin staining, primary and secondary antibody staining were omitted and a diluted solution of phalloidin AlexaFluor 488 conjugate in PBS containing diluted HCS CellMask Blue was applied for 15 min, followed by three PBS washes.

After the final staining step, the coverslips were inverted onto microscope slides (Fisher Scientific 12-550-143) over a drop of Fluoromount G (SouthernBioTech, 0100-20). Hardening was allowed overnight at room temperature. A Zeiss Axio Observer Z1 microscope was used for epifluorescence imaging using a Plan-Neofluar 40×/1.30 Oil DIC objective through Zeiss Immersol 518 F immersion oil (n_e_ = 1.518 (+23 °C)) at ambient temperature. Images were acquired blinded for construct using identical illumination and acquisition settings.

*Image analysis –* Regions-of-interest (ROI) were defined in Fiji [63] around cells using pixel intensity thresholding and manual drawing, and fluorescence intensity values were extracted from images. Binary images of ROI masks were generated in Fiji and used to calculate Pearson correlation coefficients in MatLab R2021b. Statistical analyses were performed using GraphPad Prism.

## Results

### Purification and characterisation of recombinant anti-CNPase nanobodies

Five anti-CNPase Nbs – 7E, 5E, 8C, 8D, and 10E – were raised against CNPase *via* alpaca immunisation with rat synaptosomes [26]. Alignment of the NbCNP sequences highlights the large size of their CDR3 loops (Figure 1A). Furthermore, cysteine residues are present in the CDR2s of NbCNP 5E, 8C, and 10E and in the CDR3 of 5E and 8C. All NbCNPs could be expressed in *E. coli* and purified for structural studies of CNPase-Nb complexes. DSF was used to assess NbCNP thermal stability (Figure 1B). NbCNP 7E and 10E had T_m_ of +50 °C, and NbCNP 8D was slightly more thermostable, with a T_m_ of +55 °C. Unfolding transitions were not observed for Nbs 5E and 8C, and the initial fluorescence signal resembled that of an unfolded state. As these Nbs showed binding to CNPase, this could indicate the presence of hydrophobic regions, perhaps in the CDRs, binding the assay dye in the folded state. Pull-down assays indicated that all five NbCNPs bound to the C-terminal catalytic domain of CNPase (Figure 1C).

**Figure 1.**
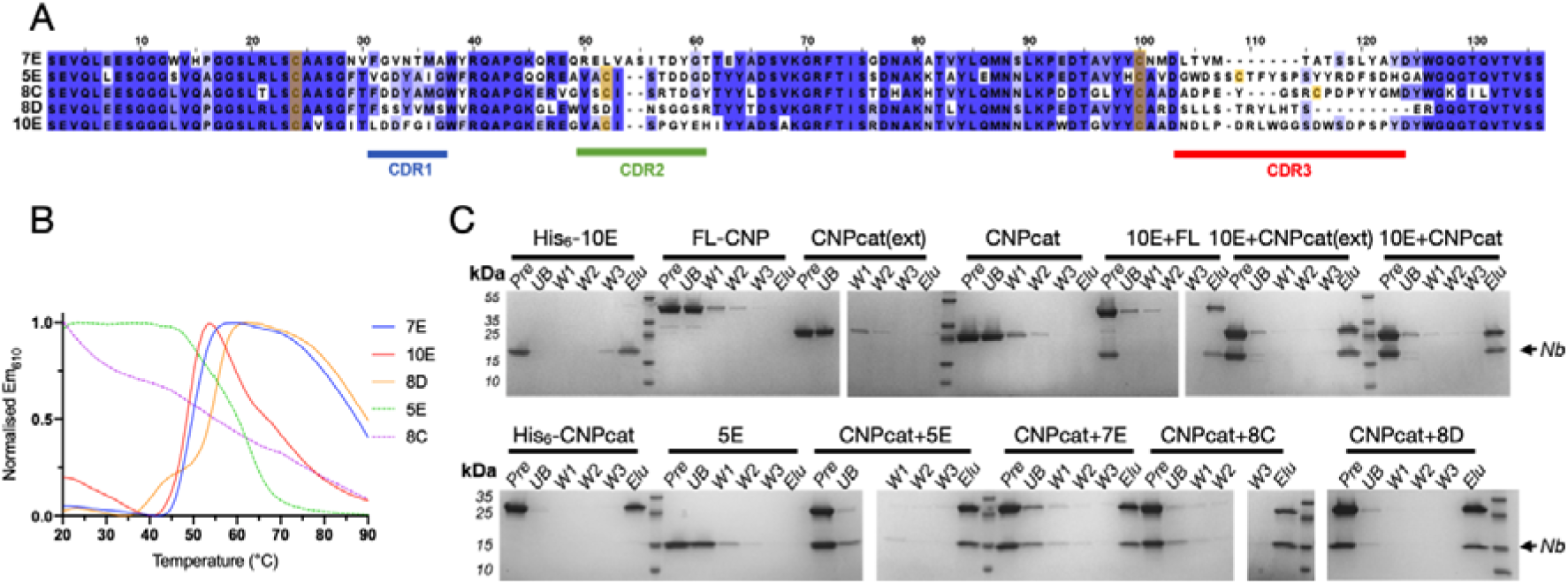
Characteristics of anti-CNPase nanobodies. A. Sequence alignment of the five NbCNPs. Cys residues are indicated in yellow. B. DSF analysis of Nb stability. C. Pulldown assay indicates all Nbs bind to the CNPase catalytic domain. FL-CNP, full-length CNPase; CNPcat, CNPase catalytic domain; CNPcat(ext), CNPase catalytic domain with C-terminal tail. Pre, sample loaded onto affinity matrix; UB, unbound fraction; W1-3, wash fractions; Elu, eluted fraction.

Mass spectrometry was used to confirm the identity of the recombinant NbCNPs (Supplementary Table 1). The observed masses were used to estimate the oxidation state of cysteines. Apparently, the conserved disulphide of both NbCNP-7E and 10E was reduced; its formation is, therefore, not essential for folding. The conserved central disulphide and the additional cysteines in the CDR2 and CDR3 of NbCNPs 5E and 8C were oxidised upon folding. The mass of 8C indicated two disulphides, and 5E gave two peaks of masses corresponding to one and two disulphides. These data suggest that Cys residues in the CDR loops of NbCNPs 5E and 8C may stabilise the paratope conformation through disulphide bridges.

### Nanobodies bind CNPase at high affinity and increase its thermal stability

The effects of all five NbCNPs on the thermal stability of full-length mouse CNPase were studied using DSF (Figure 2A, Table 3). All Nbs showed thermal stabilisation of CNPase, with the strongest stabilisation observed for NbCNPs 5E and 8D. The reproducibility of the observation was shown by an additional label-free nanoDSF experiment for the CNPase-8D complex (Figure 2B, Table 3).

**Figure 2.**
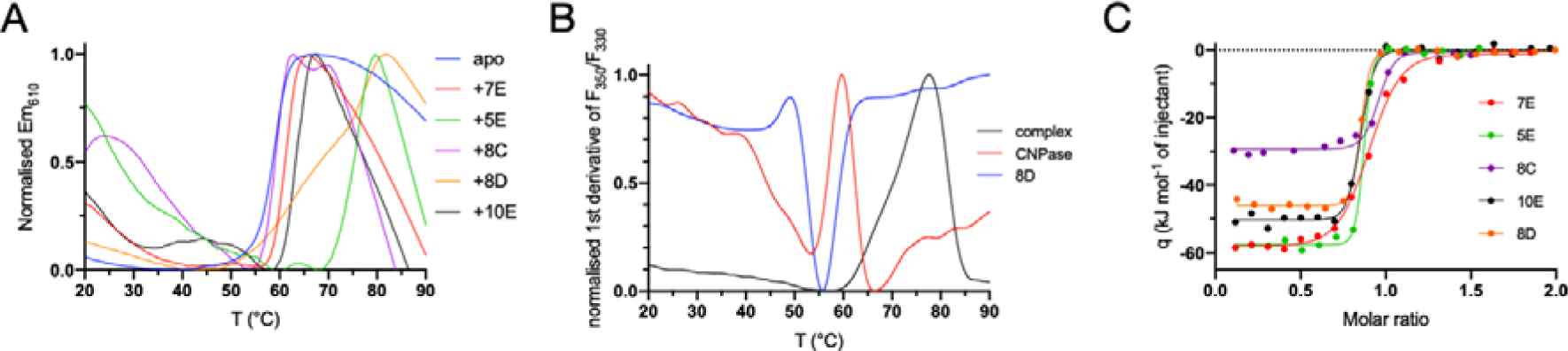
NbCNP binding to full-length mCNPase. A. Interaction probed via a DSF thermal shift assay. B. nanoDSF experiment for nanobody 8D. C. ITC titration of Nbs into CNPase.

**Table 3.** Thermal stabilisation of full-length CNPase upon NbCNP binding. T _m_ values were determined using DSF (N=3). Nanobody 8D was additionally studied using label-free nanoDSF.

| sample | $T_m$ (°C) | $\Delta T_m$ (°C) |
| --- | --- | --- |
| CNPase |  |  |
| DSF | 59.2 ± 0.1 | - |
| nanoDSF | 59.7 ± 0.03 | - |
| +7E | 61.3 ± 0.3 | +2.1 |
| +5E | 76.2 ± 0.3 | +17.0 |
| +8C | 68.4 ± 0.5 | +9.2 |
| +8D |  |  |
| DSF | 77.2 ± 0.3 | +18.0 |
| nanoDSF | 77.4 ± 0.1 | +17.7 |
| +10E | 63.4 ± 0.6 | +4.2 |

The binding affinity was examined using ITC (Figure 2C, Table 4). Most of the NbCNPs bound with a *K*_d_ of 0.5-15 nM; NbCNP 7E showed affinity decreased by an order of magnitude. The contributions of the enthalpy and entropy terms varied, suggesting diverse modes of enthalpy-driven binding.

**Table 4.** Thermodynamic parameters of NbCNP binding to CNPase catalytic domain. Error margins are from data fits for single experiments.

| Nb | Number of sites | $K_d$ (nM) | $\Delta H$ (kJ mol<sup>-1</sup>) | -T$\Delta S$ (kJ mol<sup>-1</sup>) | $\Delta G$ (kJ mol<sup>-1</sup>) |
| --- | --- | --- | --- | --- | --- |
| 5E | 0.820 ± 0.003 | 2.2 ± 0.8 | -57.70 ± 0.42 | 8.27 | -49.43 |
| 7E | 0.863 ± 0.005 | 83.3 ± 9.3 | -59.25 ± 0.52 | 18.78 | -40.47 |
| 8C | 0.914 ± 0.007 | 16.6 ± 5.5 | -29.59 ± 0.52 | -14.82 | -44.41 |
| 10E | 0.804 ± 0.005 | 9.8 ± 2.9 | -50.63 ± 0.64 | 4.89 | -45.74 |
| MaBP-10E | 0.836 ± 0.004 | 13.4 ± 3.1 | -75.60 ± 0.79 | 31.01 | -44.90 |
| MaBP-8D | 0.837 ± 0.004 | 0.2 ± 0.4 | -45.90 ± 0.49 | -9.43 | -55.35 |
| 8D | 0.800 ± 0.002 | 2.5 ± 1.6 | -46.11 ± 0.36 | -3.02 | -49.13 |

Initially, insufficient amounts of pure NbCNP-8D could be obtained for the ITC assay, and the NbCNP 8D fusion with the intact His_6_-MaBP (maltose binding protein) was used for the ITC experiment. As a control, the binding of the 10E nanobody with and without the MaBP tag was studied. The MaBP-8D fusion showed the highest binding affinity of all, with a *K*_d_ <1 nM. MaBP-10E bound with a *K*_d_ similar to NbCNP 10E with the fusion tag removed, and therefore, the *K*_d_ measured for MaBP-8D likely approximates to that of the untagged Nb, and MaBP has no effect on binding. An experiment carried out on purified Nb 8D confirmed that 5E and 8D were the NbCNPs with the highest affinity towards the mCNPase catalytic domain; in fact, the affinities are so high that they cannot be accurately determined using ITC under the current conditions. Altogether, ITC demonstrated that all NbCNPs bind the catalytic domain of mCNPase with high affinity, with varying degrees of enthalpy-entropy compensation. However, the Nbs may additionally bind the N-terminal PNK-like domain in the full-length protein, which will be a subject for future experiments. The result additionally shows that a fusion of NbCNPs with a larger partner, MaBP, has no effect on binding affinity.

#### Structure of CNPase-Nb complexes

The structures of the CNPase catalytic domain in complex with all five NbCNPs were determined by X-ray crystallography (Figure 3, Table 2). In all cases, the bound Nb could be almost completely built into the electron density, but flexibility in some solvent-exposed loops of CNPase resulted in poor density. Each NbCNP has a different epitope on the CNPase catalytic domain (Figure 3A). Nbs 8C and 8D bind the furthest away from the active site (Figure 3A,E,F).

**Figure 3.**
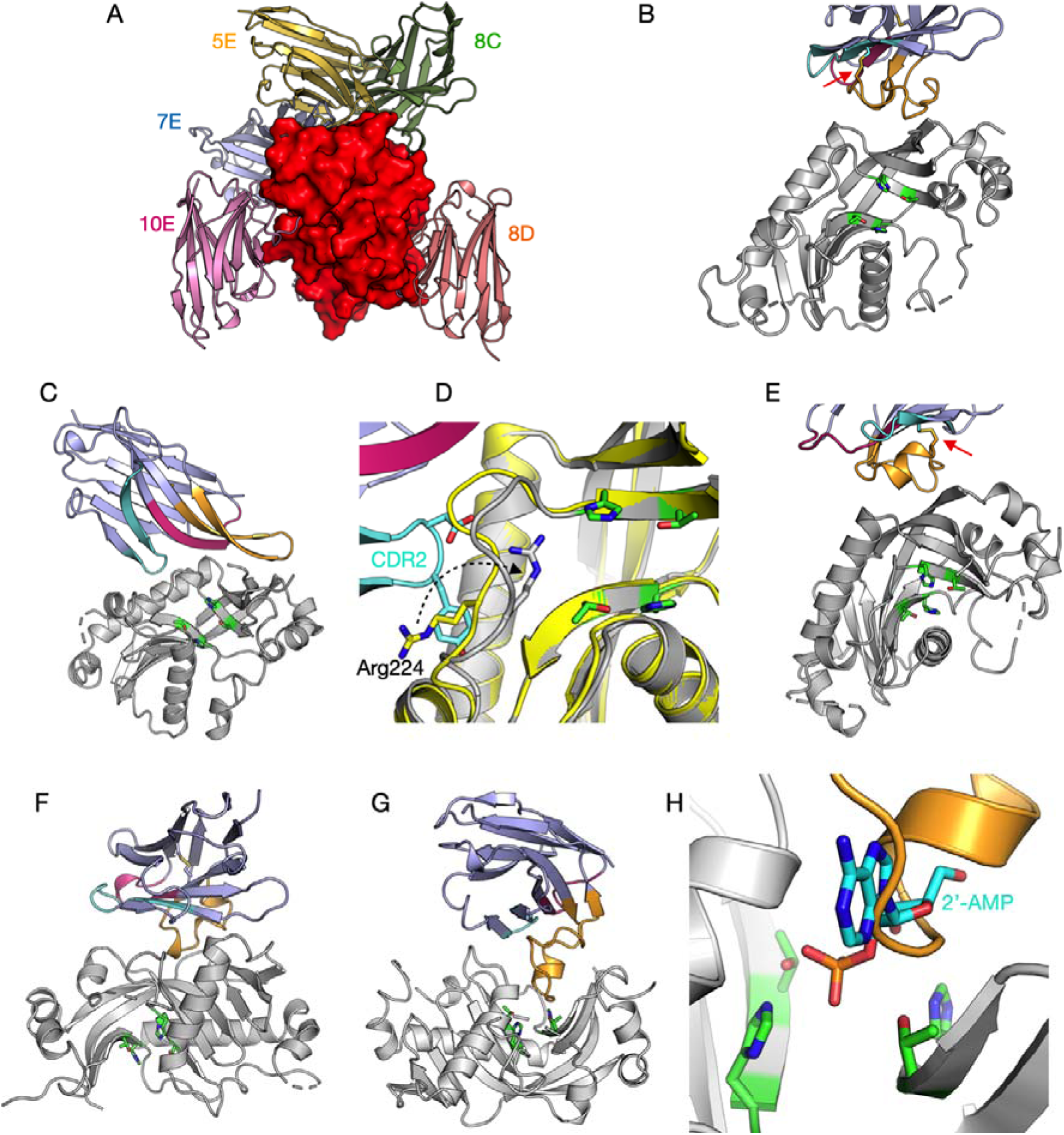
The 5 anti-CNPase nanobodies all have different epitopes. A. The individual crystal structures have been superimposed based on the CNPase catalytic domain (red surface). Individual structures: B. 5E. C. 7E. D. Zoom into the conformational change of Arg224 (arrow) near the active site, induced by the CDR2 loop. E. 8C. F. 8D. G. 10E. H. Zoom into the active site in the 10E complex, showing CDR3 overlapping with the binding mode of the reaction product 2’-AMP (cyan) [16]. The CDR loops (CDR1, magenta; CDR2, cyan; CDR3, orange) and catalytic residues (green; two HxT motifs) are highlighted in panels B-H. The disulphide bridge between CDR2 and CDR3 in nanobodies 5E and 8C is indicated with a red arrow (panels B,E).

In NbCNPs 5E and 8C, the CDR3 loop is linked to the central Nb fold *via* an extra disulphide bond between CDR2 and CDR3 (Figure 3B,E), as supported by the mass spectrometry data (Supplementary Table 1). Accordingly, the CDR3 forms a large flat paratope, predominantly binding a hydrophobic surface patch on CNPase. Both 5E and 8C interactions are primarily mediated by aromatic residues concentrated in the CDR3.

In contrast, 7E and 10E lack a cysteine in the CDR3, which extends away from the central Nb fold to bind the epitope. In 7E, the CDR1 and CDR2 bound to CNPase in proximity to the active site altering the conformation of a flexible loop and Arg224, while the elongated CDR3 loop formed β-sheet-like interactions with the CNPase C-terminal region, which lies close to the N terminus of the catalytic domain (Figure 3C,D). A large portion of the 7E CDR3 loop does not bind CNPase, which can explain its lower affinity and suggests that the loop might bind to the interdomain interface in full-length CNPase.

The elongated CDR3 of NbCNP-10E extends into the CNPase active site and overlaps with the substrate/product binding site (Figure 3G,H). Moreover, the CDR1 and CDR2 of 10E bound a second CNPase molecule in the crystal, apparently fixing the protein in a homodimeric state in the crystal. Possible dimerisation was confirmed using PISA, and the calculated ΔG^int^ of -64.02 kJ mol^-1^ suggested a stable assembly. This putative dimer interface formed between hydrophobic surface patches of CNPase and buried 930 Å^2^ of solvent-accessible surface of each monomer. This dimerisation, also suggested by the SAXS experiments below, could represent an oligomeric state of full-length CNPase *in vivo*, since it was picked up by the 10E nanobody. In addition, it may be linked to the inhibitory mode of nanobody 10E.

#### Solution structure of CNPase catalytic domain-nanobody complexes

To observe and validate the complexes in solution, we conducted synchrotron SEC-SAXS experiments on the CNPase catalytic domain and the NbCNPs (Figure 4, Table 5). All CNPase catalytic domain-Nb complexes eluted as single peaks, suggesting one dominant oligomeric state. All complexes were 1:1 heterodimers and fit well to the SAXS data and closely resembled *ab initio* chain-like models (Figure 4, Supplementary Figure 1), except for the NbCNP10E, for which the complex had a slightly higher apparent molecular weight. This may indicate Nb-induced dimerisation of the CNPase catalytic domain or *vice versa*, which could relate to the mode of inhibition of CNPase catalysis by 10E. The assembly from the crystal that fit the SAXS data best contained 2 molecules of CNPase and one 10E molecule. For the case of NbCNP 10E, as well as full-length CNPase, additional studies will be required to conclude on detailed structural properties of the complex.

**Figure 4.**
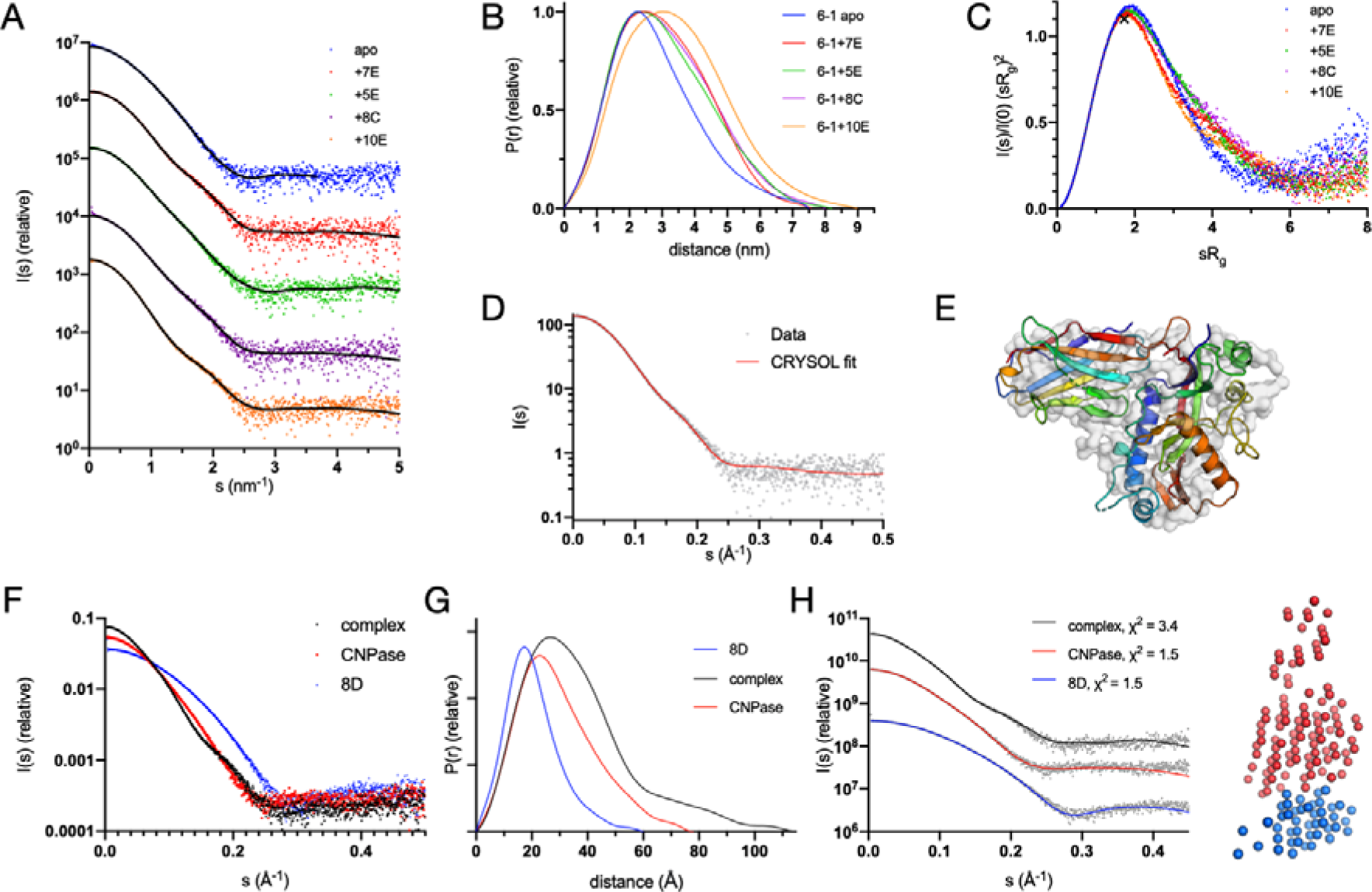
Conformation of CNPase-nanobody complexes in solution. A. Scattering data, with fits of chain-like ab initio models as lines. B. Distance distribution. C. Dimensionless Kratky plot. The position of the cross corresponds to a theoretical maximum for a globular particle. D. Fit of the CNPase-NbCNP 7E crystal structure to the raw SAXS data. E. Superposition of the chain-like SAXS model of CNPase-NbCNP 7E (surface) and the crystal structure (cartoon). Fits for all complexes to the SAXS data are shown in Supplementary Figure 1. F. Scattering data for the NbCNP 8D complex. G. Distance distribution for NbCNP 8D. H. Fit of the multi-phase MONSA model to all datasets simultaneously. The model of the complex is shown on the right (red – CNPase, blue – NbCNP 8D). The NbCNP 8D complex data are shown separately due to the different expression constructs used for nanobody purification and for comparing all components in the same experiment.

**Table 5.**
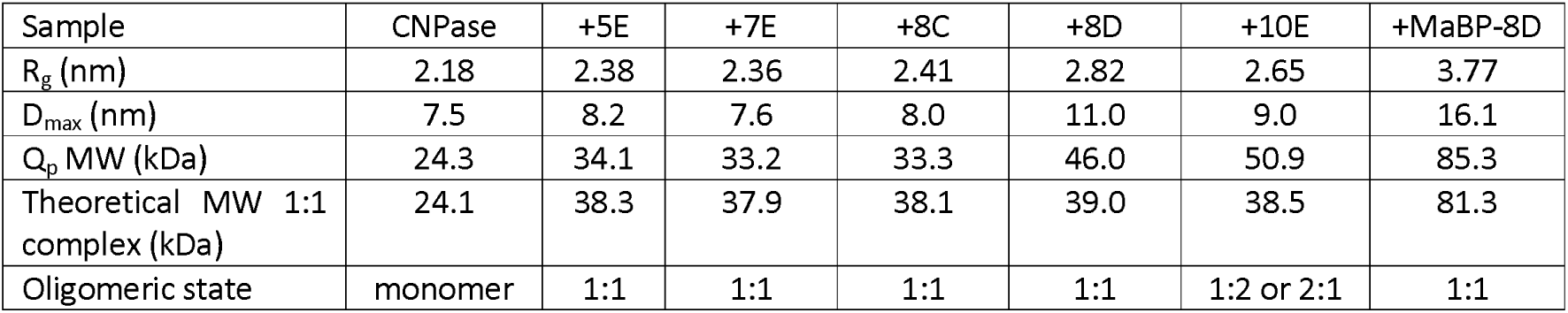
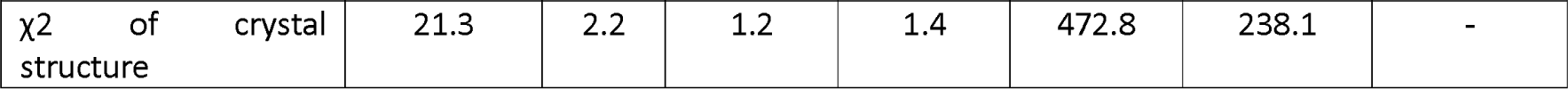
Size and shape parameters of CNPase-NbCNP complexes derived from SAXS data. The catalytic domain of mCNPase was used in the experiments. The fits of the crystal structures (from CRYSOL) to the solution SAXS data indicate that CNPase alone is more extended in solution than in the crystal, the CNPase-8D complex in solution has an extended tail, and the CNPase-10E complex stoichiometry differs from 1:1 (see text and Supplementary Figure 1 for details).

The varying R_g_ and D_max_ among the complexes suggested different binding modes. The NbCNP 8D complex with CNPase shows a higher D_max_ due to the different construct used to produce the Nb for these experiments, with a short, extended tail. The NbCNP 8D SAXS data alone showed the same elongated feature (R_g_ 1.57 nm, D_max_ 6.0 nm), larger than expected for a compact Nb single domain. This explains why the crystal structure provides a poor fit to the experimental SAXS data in this case (Table 5, Supplementary Figure 1). As an alternative modelling protocol, a two-phase model for the CNPase-8D complex was built, using data for both components and the complex simultaneously (Figure 4F-H). The model fits both the individual components and the complex, supporting a 1:1 complex between CNPase and NbCNP 8D. The slightly poor fit of the crystal structure to the raw SAXS data in the case of NbCNP 8D is likely explained by disordered regions not ordered in the crystal, but present in solution. The stoichiometry was additionally confirmed by an experiment using the MaBP-NbCNP 8D fusion bound to CNPase (Table 5).

#### CNPase activity

To explore the effect of Nb binding on CNPase function, the enzymatic activity of the full-length mCNPase was measured in complex with the nanobodies NbCNP-7E, -5E, and -10E (Figure 5, Table 6). Full-length mCNPase-nanobody complexes purified by SEC were used in the experiment. NbCNP-5E had little to no effect on CNPase activity (Table 6). However, with NbCNP-7E, a slight increase in k_cat_ was apparent, indicating activation of CNPase upon 7E binding close to the active site. NbCNP-10E was a non-competitive inhibitor of CNPase (Figure 5A,B), facilitating a considerable reduction in k_cat_, with an unchanged K_M_. The inhibition was potent, with an apparent K_i_ of 0.6 nM (Figure 5C, Table 6), in line with the high affinity of the nanobody towards CNPase. Non-competitive inhibition results from inhibitor binding to both the free enzyme and the enzyme-substrate complex; confirmation of this inhibition mode will require experiments with a wider range of nanobody concentrations in the future.

**Figure 5.**
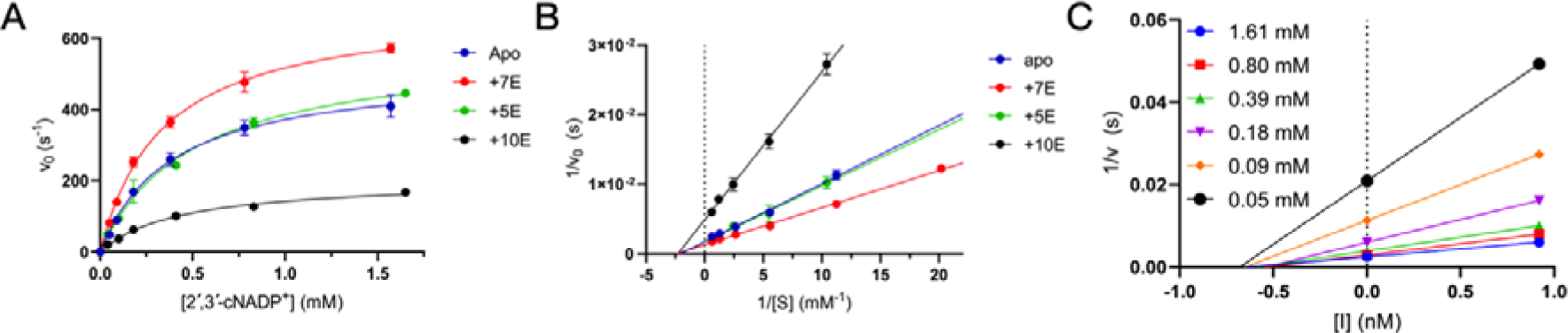
CNPase catalytic activity in the presence of selected nanobodies. A. Initial rates of 2’,3’-cNADP^+^ hydrolysis. B. Lineweaver-Burk plot indicates non-competitive inhibition by nanobody 10E. C. Non-competitive Dixon plot in the presence of nanobody 10E. [I] denotes the concentration of 10E, and the x-axis intercept gives apparent -K_i_.

**Table 6.**
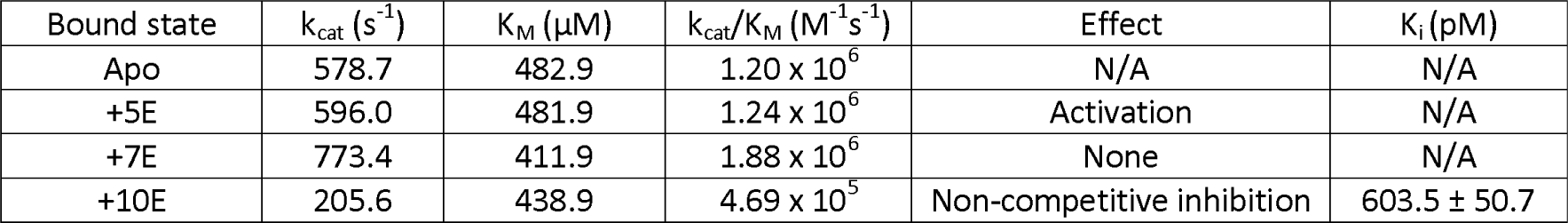
Kinetic parameters of full-length mCNPase in the presence of selected nanobodies. N/A, not applicable.

| Bound state | $k_{\text{cat}}$ (s<sup>-1</sup>) | $K_M$ (μM) | $k_{\text{cat}}/K_M$ (M<sup>-1</sup>s<sup>-1</sup>) | Effect | $K_i$ (pM) |
| --- | --- | --- | --- | --- | --- |
| Apo | 578.7 | 482.9 | $1.20 \times 10^6$ | N/A | N/A |
| +5E | 596.0 | 481.9 | $1.24 \times 10^6$ | Activation | N/A |
| +7E | 773.4 | 411.9 | $1.88 \times 10^6$ | None | N/A |
| +10E | 205.6 | 438.9 | $4.69 \times 10^5$ | Non-competitive inhibition | $603.5 \pm 50.7$ |

### Nanobody specificity in imaging applications

NbCNP 5E and 8D were selected for further validation because of their strong affinity towards the catalytic domain of mCNPase. NbCNP 10E was chosen because of its specific binding into the active site. NbCNPs 5E, 8D, and 10E with an ectopic cysteine at the C-terminus were prepared, and site-specific conjugation was performed using maleimide-functionalised fluorophores [25, 26].

For a specificity test, we prepared teased sciatic nerves from wild-type mice and mice lacking CNPase [8] and stained them with the fluorescently labelled NbCNPs. We obtained specific signals on the wild-type nerves with all three selected nanobodies (Figure 6A), while no signal was observed from the CNPase-deficient tissue (Figure 6B), indicating specificity of the signal. All nanobodies detected mCNPase throughout the non-compacted regions of the myelin sheath; a stronger signal can be observed on the paranodes where the nanobodies have better access to target CNPase.

**Figure 6.**
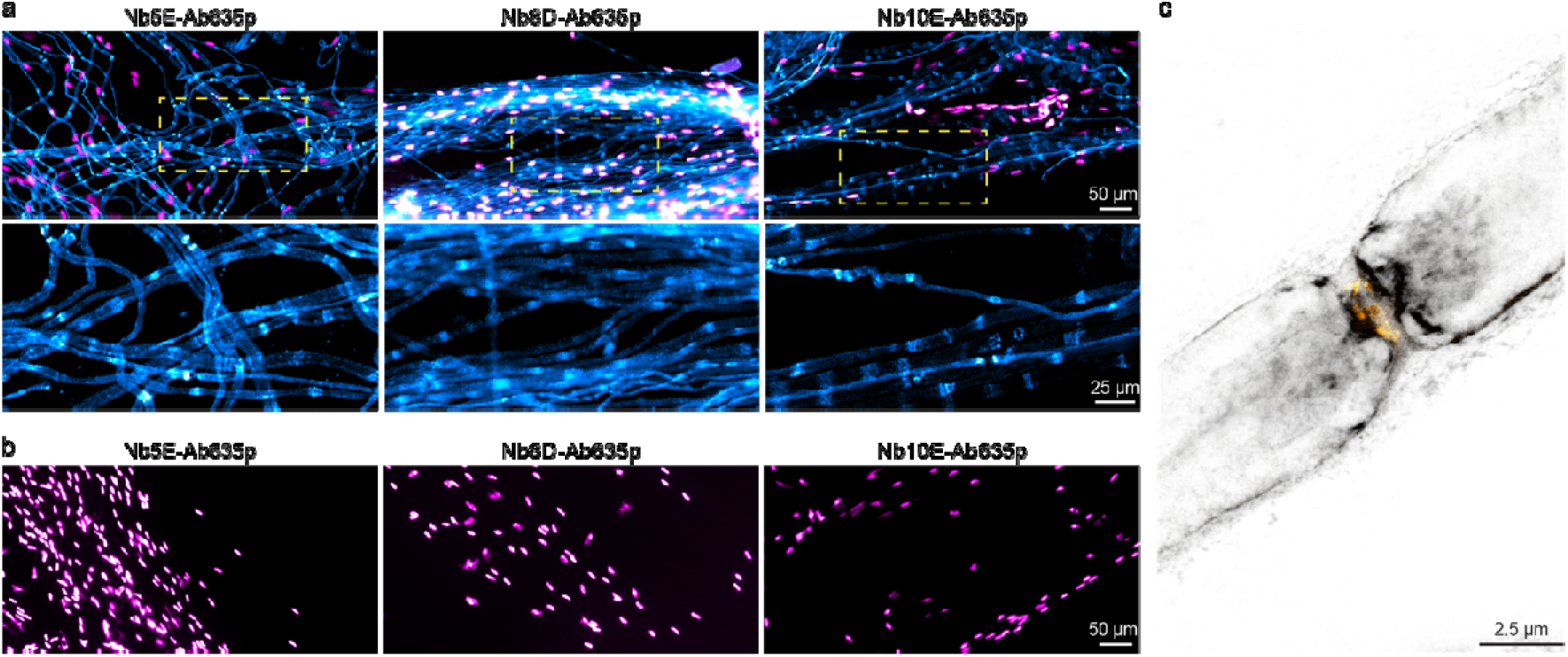
Specific staining of mCNPase using fluorescently labelled nanobodies. (a) Epifluorescence images of teased sciatic nerves from wild-type mice stained with Nb5E, Nb8D, and Nb10E displayed in cyan, nuclei stained with DAPI displayed in magenta. (b) Teased sciatic nerves from mice lacking CNPase display no staining (cyan) and nuclei in magenta. (c) 2-color STED image using Nb8D-Ab635p (grey) and voltage-gated sodium channels (Nav1.6) (yellow).

We then performed 2-color STED microscopy using Nb8D-Ab635p and antibodies against voltage-gated sodium channels (Nav1.6) on the node of Ranvier, confirming that the nanobody signal is enhanced on the paranodes (Figure 6C).

Finally, we used the NbCNPs for a challenging test by performing immunohistochemistry on mouse brain tissue sections (Figure 7). We observed a highly efficient staining of myelinated tracts in the brain, and there was virtually no background staining in regions where myelin was not expected. In addition, we could observe the different affinities of NbCNPs by comparing the fluorescence intensities, for which NbCNP 8D was significantly brighter, providing an excellent signal-to-noise ratio. Based on the crystal structures, these three nanobodies (5E, 8D, 10E) could eventually be used in an oligomeric cocktail to enhance the signal further. The crystal structures indicate that the nanobodies should not sterically interfere with the binding of one another, as they bind onto different faces of CNPase.

**Figure 7.**
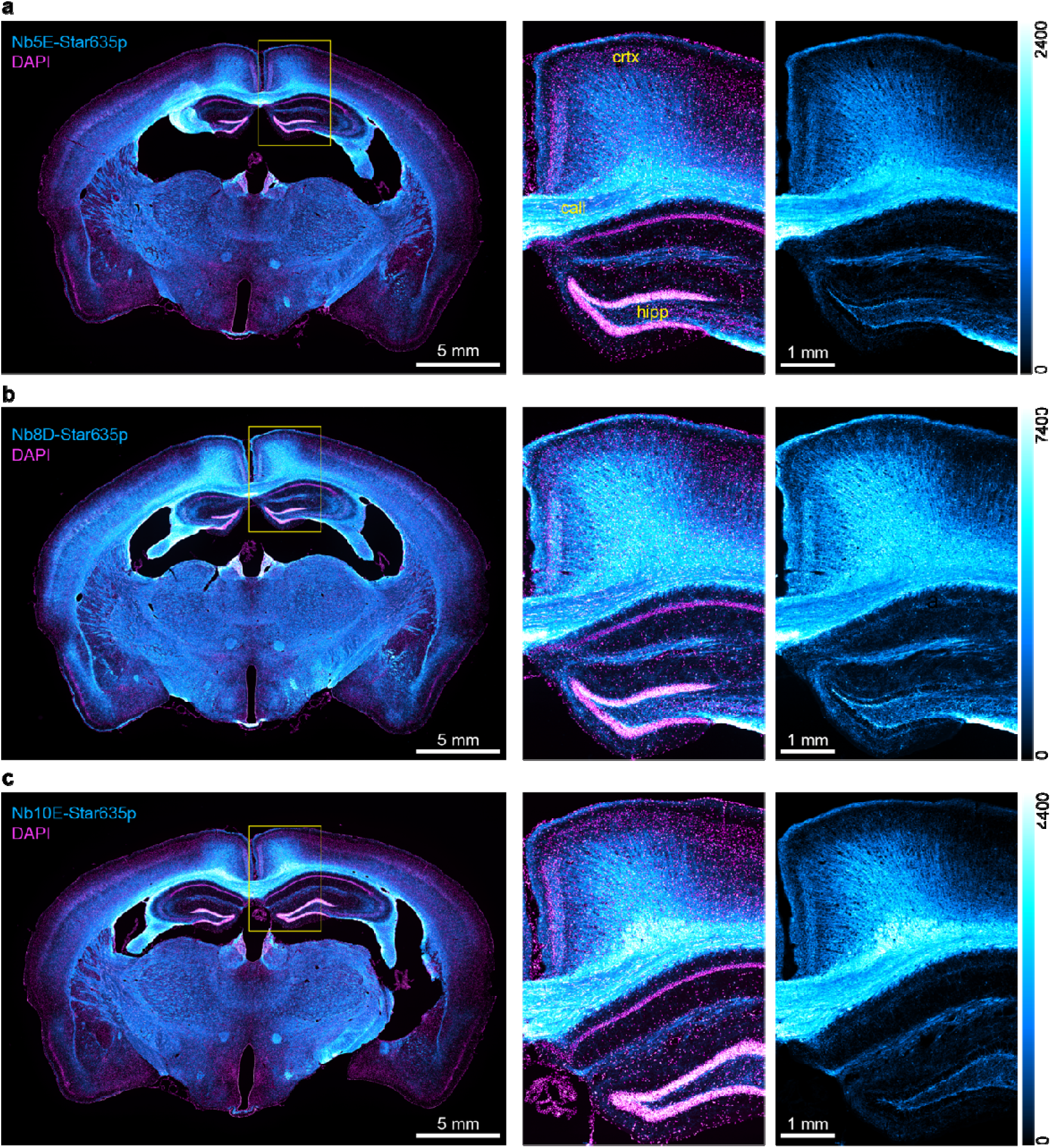
Immunohistochemistry on mouse brain tissue sections. Epifluorescence images of coronal mouse brain sections. In cyan, the stainings of (a) Nb5E, (b) Nb8D, and (c) Nb10E. Nuclei stained with DAPI in magenta. Yellow rectangles show the zoomed region, including the hippocampus (hipp), corpus callosum (call), and cortex (crtx) as an overlay (CNPase + nuclei) and single color (CNPase). The intensity scale for the cyan look-up table is displayed to the right of each panel.

### Expression of NbCNPs as intrabodies (iNbCNPs)

As a means to validate the selected NbCNPs as intrabodies, the nanobodies 5E, 8D, and 10E fused to mRuby3 were co-expressed with TOM70-EGFP-CNPase (catalytic domain) in COS-7 cells. These nanobodies were selected due to their high affinity and effect on CNPase activity. In this system, an EGFP-tagged CNPase is anchored to the mitochondrial outer membrane (TOM70), and if Nbs are functional as iNbs in the reducing cytoplasmic environment of living cells, we expected a colocalisation between EGFP and mRuby3 signals. All three tested intrabodies showed strong co-localisation (Pearsońs correlation) with the CNPase anchored to mitochondria, while expression of mRuby3 alone as negative control shows poor specific colocalisation, demonstrating that these NbCNPs can be used as iNbs against CNPase (Figure 8). Especially interesting for future applications is the NbCNP 10E, which also inhibits CNPase catalytic activity (Figure 5).

**Figure 8.**
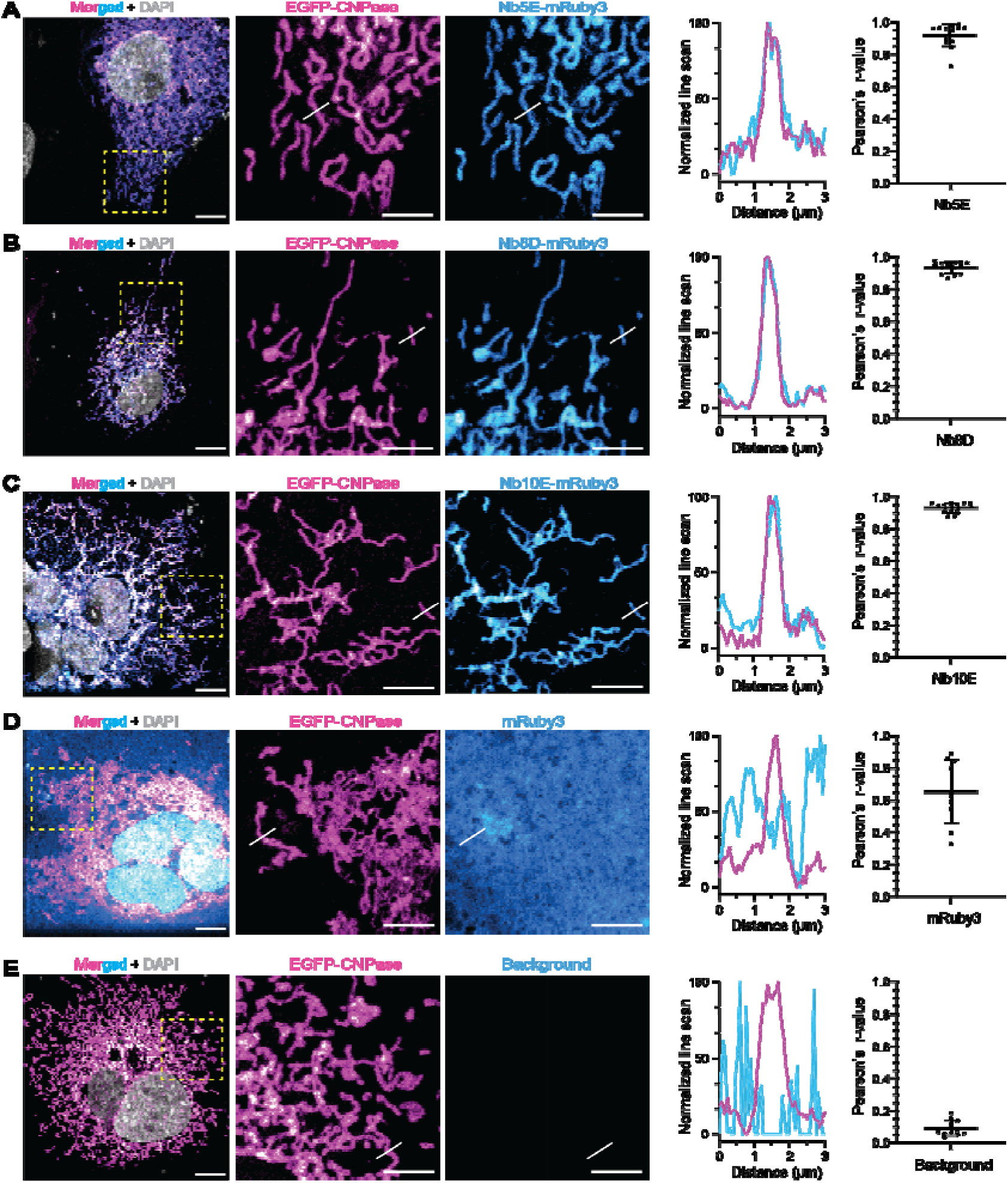
Nanobodies as intrabodies in COS-7 cells. Confocal images were acquired after COS-7 cells were co-transfected with TOM70-EGFP-CNPase and an intrabody fused to mRuby3. (a) Merged signals in a wide field of view, followed by zoomed regions (yellow square) displaying the mitochondria structures by TOM70-EGFP-CNPase (magenta) and the signal from mRuby3 fused to iNbCNP 5E. White lines denote the position where the line-intensity profile (line-profile) is shown in the graph. Pearsońs correlation between the mitochondrial EGFP signal (magenta) and the mRuby3 signal from the intrabody (cyan) was calculated from 9 independent cells. The graph represents the mean ± standard deviation. The same analysis was performed for iNbCNP 8D ( **B**), iNbCNP 10E ( **C**), and the controls with cytosolic mRuby3 not fused to any intrabody ( **D**), or no intrabody transfected ( **E**). This last control rules out potential signal bleed-through from EGFP into the mRuby3 channel.

All five nanobodies were then tested for expression as intrabodies in cultured oligodendrocytes (Figure 9, Supplementary Figure 2), which is a much more complex cellular system. The iNbCNPs 7E, 8D and 10E, but not 5E and 8C, were expressed in cultured oligodendrocytes and co-localise with CNPase immunostaining in cytoplasmic channels, as well as in cytoplasmic patches amongst compaction zones, at day 5 of differentiation. As expected, there was little to no colocalisation with myelin basic protein (MBP) [64]. Since CNPase interaction with actin filaments has been shown [20], actin was stained with phalloidin. Some potential iNbCNP co-localisation was detected with actin, although most actin at day 5 of differentiation is present as a rim at the edges of the cell. Currently, we do not know if the binding of iNbCNPs to CNPase affects its ability to bind actin filaments in living cells, but at least some of them likely do.

**Figure 9.**
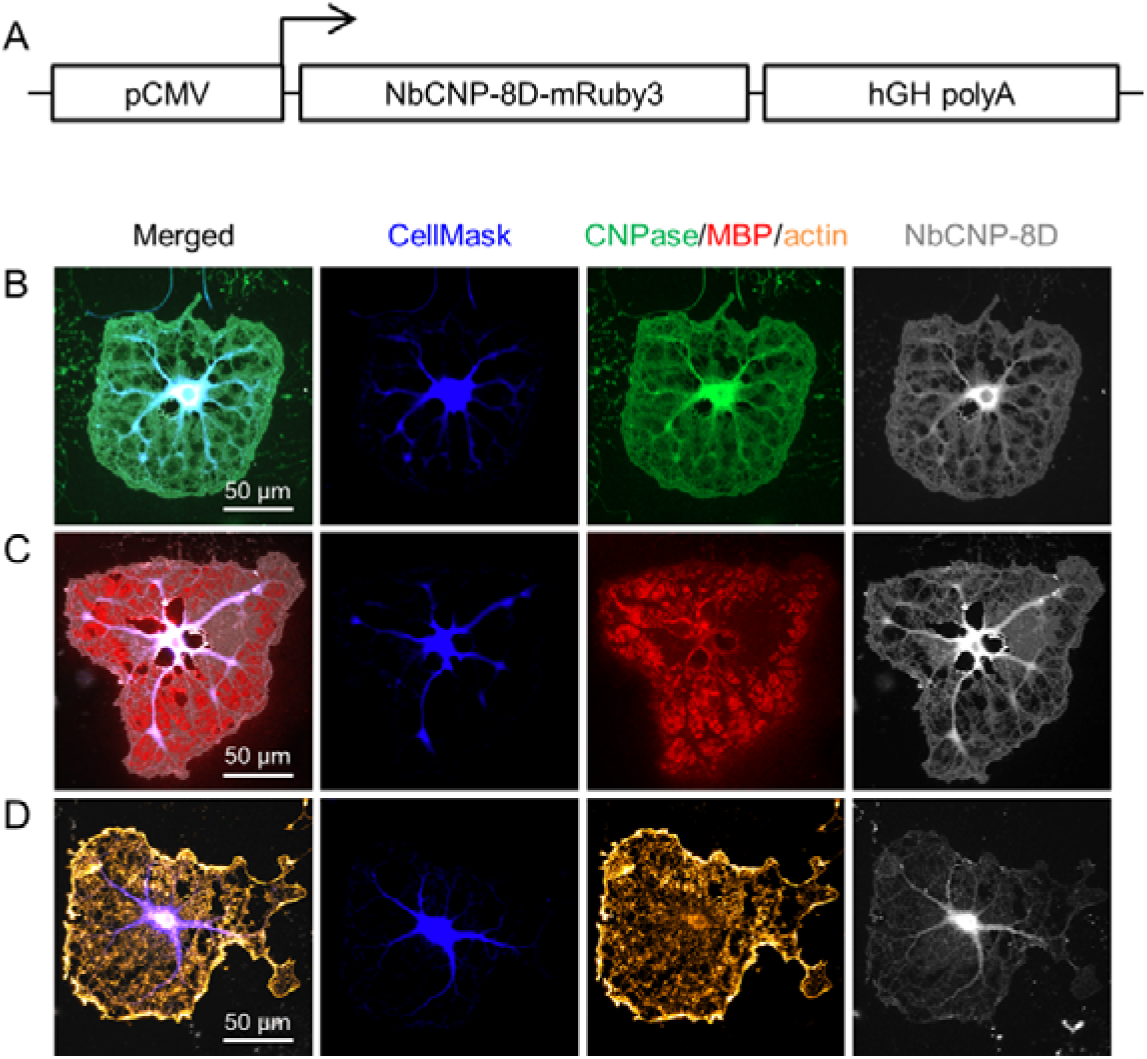
Expression of NbCNPs as intrabodies in cultured oligodendrocytes. Shown is the experiment with iNbCNP 8D. Experiments with the other nanobodies are shown in Supplementary Figure 2. A. The iNbCNP 8D construct architecture, with fused mRuby3, used in transfection. NbCNP-8D-mRuby3-expressing cells were stained for either CNPase (B), MBP (C), or actin (D). Fluorescence for the labelled iNbCNP was directly recorded (gray).

The iNbCNP-mRuby3 fusions 5E and 8C did not produce fluorescence in the oligodendrocytes; hence, there was either no expression or protein folding/aggregation. Interestingly, these are the two nanobodies with an extra disulphide bridge linking CDR2 and CDR3, which may reflect a folding issue in the reducing cellular environment; the putatively different redox environments of COS-7 cells and oligodendrocytes likely explain some of the differences between experiments. In addition, NbCNP 10E apparently decreased CNPase staining by the conventional antibody (Supplementary Figure 2). 10E may bind the same epitope as the antibody used for immunostaining in the experiment, or oligodendrocytes might remove/degrade non-functional CNPase when inhibited by iNbCNP 10E.

## Discussion

### Nb screening and production

The anti-CNPase nanobodies presented here resulted from a fortuitous immune reaction against myelin protein contaminants in a rat brain synaptosome preparation that was used for alpaca immunisation. Since the major myelin proteins, including CNPase, are abundant proteins in nerve tissue, this was not fully unexpected. The result suggests high antigenicity of CNPase, given that it was an impurity in the preparation; interestingly, all epitopes for the nanobodies lie on the C-terminal catalytic domain, each with its distinct epitope.

With some modifications in the protocols, all nanobodies could be produced recombinantly on a large scale, and some of the unexpected yield and solubility issues might have been caused by the extra Cys residues present in the CDR loops. For two of the nanobodies, these form a disulphide bridge between CDR2 and CDR3, stabilizing the CDR conformation. Such cysteine residues in exposed loops are likely reactive and can cause problems during recombinant production on a large scale.

The high-quality recombinant anti-CNPase nanobodies can be additionally modified and optimised for further applications, which – as shown here – can include structural biology, imaging, and functional intervention. As a key goal in myelin structural biology, they are currently being used to facilitate experimental structure solutions of full-length CNPase and its molecular complexes at high resolution. In addition, in line with recent technical developments for nanobody-based tools [65–67], we will pursue the possibility of using CNPase-nanobody complexes as fiducials for cryo-EM experiments.

## Structure of the CNPase-nanobody complexes

The five NbCNPs all had different epitopes, as suggested by their varying CDR sequences. Hence, CNPase carries several potential immunogenic surfaces, which include both pockets and rather flat surfaces. Interestingly, all these epitopes were in the C-terminal phosphodiesterase domain. It is possible that CNPase in the preparation was in an oligomeric state, which could hide the N-terminal domain from presenting potential epitopes, or the N-terminal domain simply was less immunogenic under these conditions. Some of the nanobodies may also interact with the N-terminal domain in addition to the main epitope in the CTD. Our initial observation that full-length CNPase forms oligomers in the presence of the nanobodies, as well as the possible dimerisation of the catalytic domain alone in the presence of nanobody 10E, is therefore of further interest, with regard to both structure and function of CNPase.

Nanobody 10E bound directly deep into the CNPase active site and acted as an inhibitor. Based on the crystal structure and earlier structural data, the CDR3 loop should prevent substrate binding into the active site. Therefore, it was surprising that 10E was a non-competitive inhibitor instead of a competitive one. One clue to the mechanism came from SAXS data, indicating a 2:1 complex between the catalytic domain and 10E. Further high-resolution studies will be required to understand the full mechanism of inhibition, preferably in the context of full-length CNPase.

Another intriguing case was nanobody 7E. At first glance, it appeared not to interact with the active site. However, since activity assays indicated an increase in activity, the interactions of the nanobody with the Ill3-β2 loop in the vicinity of the active site appear to be functional. The conformational change of Arg224, moving much closer to the active site, may be a means to increase catalytic efficiency of CNPase, for example, by changing the electrostatic potential in the active site – or by affecting the water network, which is important for substrate recognition and catalysis [5, 17, 18]. This effect is of interest when considering the reasonably well-defined catalytic mechanism of CNPase [6, 16–18, 68] and its implications. For example, the mutation P225G in the neighbouring residue in the same loop caused a similar effect on CNPase catalytic properties as 7E [18]; therefore, the Ill3-β2 loop apparently plays a role in fine-tuning the enzymatic reaction of CNPase. Similarly, Arg307 on the opposite side of the active-site cavity was shown to be important for catalysis; its mutation to Gln caused an increase in both k_cat_ and K_M_ [18].

Nanobodies 5E, 8C, and 8D had no apparent effects on CNPase catalytic activity in the current study, when 2’,3’-cNAPD^+^ was used as substrate. They might, however, affect activity towards larger substrates, such as RNA. Of note, all five CNPase-nanobody complexes were submitted for structure prediction in CASP15, in order to explore the ease of predicting nanobody-target interactions and correct paratope/epitope conformations. As described [69], one of them (complex with Nb 8C) could not be predicted correctly by any of the contestants. This is likely linked to the conformation of the CDR3 loop in the 8C structure, which contains a short helix held in place by a disulphide bridge.

### Anti-CNPase nanobodies as tools for imaging and functional intrabodies

CNPase is one of the most used markers for myelinating glia in the CNS and PNS [70–73]. It is localised in non-compact myelin areas, including cytoplasmic channels, paranodal loops, and Schmidt-Lanterman incisures [20, 74]. Hence, we see an interest and potential in further developing the anti-CNPase nanobodies into versatile tools for both routine and high-resolution imaging applications and functional studies.

Imaging of myelin is a standard tool to visualise the structure of the nervous system. We have shown here that the anti-CNPase nanobodies are highly specific; no signal was visible by staining CNPase-deficient nerves. Direct labelling of Nbs allows a simplified single-step immunostaining, and their small size combined with their strong affinity provides high signal-to-noise, allowing clean staining on teased nerves and fast penetration on whole brain tissue sections. The strong signal and small linkage error make these Nbs ideal for molecular imaging using super-resolution microscopy (*e.g.*, STED). As such, the current set of NbCNPs can already function as a toolkit for high-resolution imaging of myelinated tissue. One development option would be to combine these nanobodies simultaneously, guided by the high-resolution crystal structures, to increase the signal-to-noise even further.

Intrabodies, or antibody fragments expressed inside living cells, are gaining increasing use for functional studies of target molecules [26, 75, 76], and they are being developed towards therapeutic [77, 78] and diagnostic [79] applications. The anti-CNPase nanobodies could be used to modulate CNPase activity in living cells and to affect its interactions with natural partners, such as the actin cytoskeleton. We now have a structural basis for both inhibiting and slightly activating CNPase by nanobodies, and the wide spread of epitopes around the CNPase molecule should help in finding and optimizing nanobodies that can block molecular interactions.

## Concluding remarks

We have described the development of a panel of anti-CNPase nanobodies, which have a wide range of possible applications. Each NbCNP has a unique epitope on the C-terminal catalytic domain of CNPase. For solving the structure of full-length CNPase, they will be useful as crystallisation chaperones and/or cryo-EM fiducials. The CNPase-Nb complexes could also be developed further as a tool for the structural stabilisation of flexible proteins in structural biology. The nanobodies may provide simpler alternative tools for imaging applications, including diagnostics, where CNPase is a marker for myelinating cells. Furthermore, high-resolution imaging on myelinated tissue using advanced microscopy is possible. The expression of NbCNPs as intrabodies opens up ways of functional intervention of CNPase-related processes, modulating CNPase catalytic activity, as well as targeting functional fusion partners to defined cellular compartments.

## Acknowledgements

Early stages of this work were supported by Jane and Aatos Erkko Foundation, Finland (P.K.). F.O. was supported by Deutsche Forschungsgemeinschaft (DFG) through the SFB1286 (project Z04). DFG also supported F.O. and A.M.R. through the grant OP 261/3-1. The work was partly funded by the Momentum Career Development Programme and the Faculty of Medicine (University of Bergen) (A.R.), as well as ERASMUS+ (M.S.). We acknowledge the use of the Core Facility for Biophysics, Structural Biology, and Screening (BiSS) at the University of Bergen, which has received infrastructure funding from the Research Council of Norway (RCN) through NORCRYST (grant number 245828) and NOR–OPENSCREEN (grant number 245922). J.B.Z. was supported by NIH R01NS119823 and NIH R21NS131999-01. We thank all Zuchero lab members as well as Dr. John Vaughen and Dr. Ved Topkar at Stanford University for helpful discussions. We thank Dr. Ulrich Bergmann (Faculty of Biochemistry and Molecular Medicine, University of Oulu) for aiding in mass spectrometry and Ju Xu (University of Bergen) for technical assistance. We also thank Dr. Sandra Goebbels and Prof. Klaus-Armin Nave for discussions and support. Synchrotron beamtime and support at SOLEIL, ESRF, and DESY are gratefully acknowledged.

## Data availability

Structure factors and refined coordinates for the crystal structures are available at the Protein Data Bank, under the access codes 9ERT, 9ERU, 9ERW, 9ETL, 9ETJ.

Raw diffraction datasets are available on Zenodo, under the DOI: 10.5281/zenodo.10868891, 10.5281/zenodo.10868769, 10.5281/zenodo.10868574, 10.5281/zenodo.10868255.

All other data that support the findings of this study are available from the corresponding authors upon reasonable request.

## Supplementary files

*Supplementary Table 1. Mass spectrometric analysis of the nanobodies*

*Supplementary Figure 1. Fits of SAXS data to all crystal structures*.

*Supplementary Figure 2. Expression of all nanobodies in oligodendrocytes*.

