## Supplementary Figure 1 for "Nanobodies against the myelin enzyme CNPase as tools for structural and functional studies"

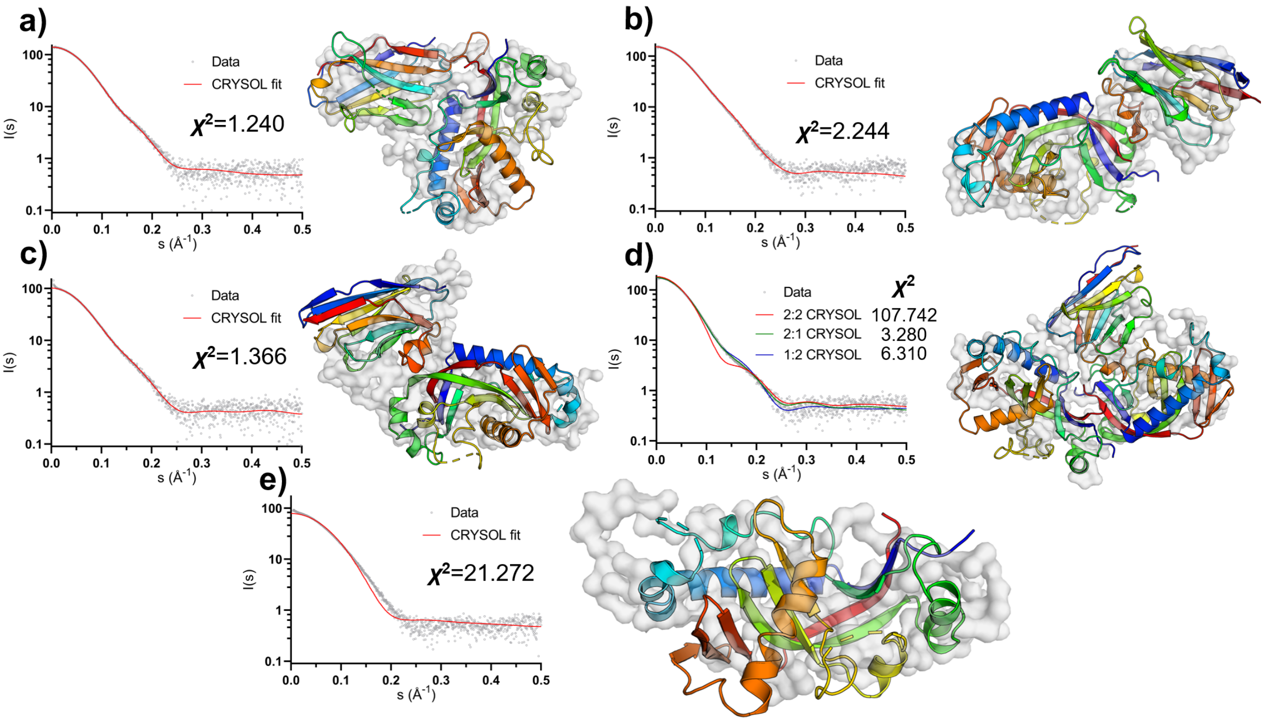


**Supplementary Figure 1.** SAXS data for complexes between mCNPase catalytic domain and NbCNPs 7E (a), 5E (b), 8C (c), and 10E (d). CNPase without Nbs is in panel (e). Shown to the right in each panel are the crystal structures (cartoons coloured in rainbow) superimposed onto *ab initio* models calculated using GASBOR (transparent grey surface). Each panel shows the $\chi^2$ of the fit, where a value close to 1.0 indicates a good fit.
