## Supplementary Figure 2 for "Nanobodies against the myelin enzyme CNPase as tools for structural and functional studies"

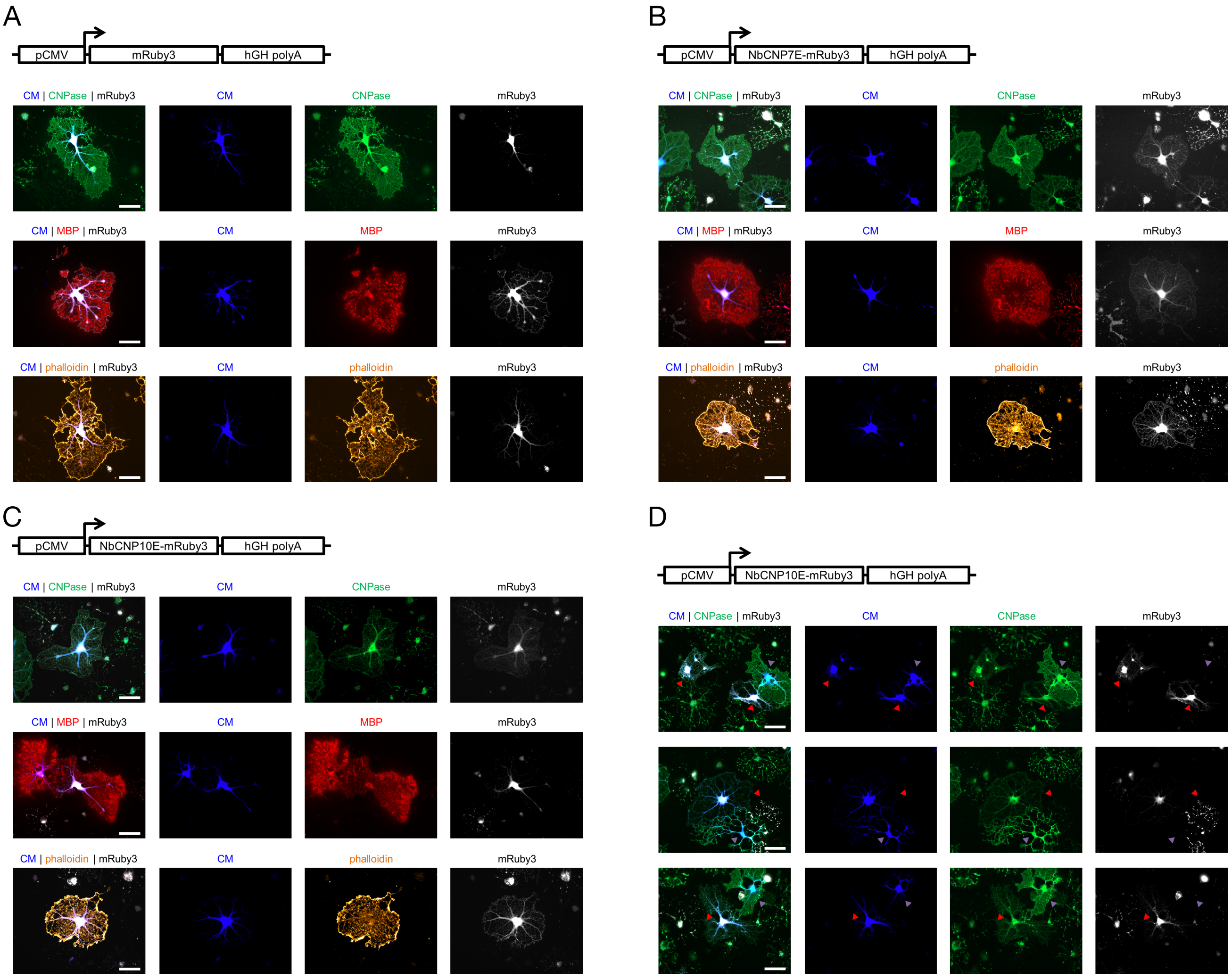


**Supplementary Figure 2.** Expression of NbCNPs as intrabodies fused to mRuby3 in cultured oligodendrocytes. A. mRuby3 alone control. B. NbCNP 7E. C. NbCNP 10E. D. NbCNP10E shows reduced staining for CNPase with conventional antibodies.
