## Supplementary Table 1 for "Nanobodies against the myelin enzyme CNPase as tools for structural and functional studies"

***Supplementary Table 1. Mass spectrometry analysis of the NbCNPs.*** *M_mi_ denotes monoisotopic mass and (S-S) a disulfide. Measurement of MaBP-8D was attempted, but the protein degraded during transport to the MS facility. The NbCNP 8D identity was confirmed by the crystal structure.*

| Protein | Calculated M_mi_ (Da) | Measured M_mi_ (Da) | Peak abundance (%) | Mass change (Da) | PTM |
| --- | --- | --- | --- | --- | --- |
| Nb 7E | 13936.59 | 13936.61  13935.62 | 66.25  29.00 | +0.02  -0.98 | None |
| Nb 5E | 14243.53 | 14241.56  14239.53 | 52.18  26.27 | -1.97  -4.00 | 1x(S-S)  2x(S-S) |
| Nb 8C | 14079.53 | 14075.51 | 94.12 | -4.02 | 2x(S-S) |
| Nb 10E | 14428.81 | 14428.83 | 99.81 | +0.02 | None |
